# Defining Structure-Function Relationships of Amphiphilic Excipients Enables Rational Design of Ultra-Stable Biopharmaceuticals

**DOI:** 10.1101/2024.06.17.599313

**Authors:** Alexander N. Prossnitz, Leslee T. Nguyen, Noah Eckman, Suraj Borkar, Samantha Tetef, Anton A. A. Auzten, Gerald G. Fuller, Eric A. Appel

## Abstract

Biopharmaceuticals are the fastest growing class of drugs in the healthcare industry, but their global reach is severely limited by their propensity for rapid aggregation. Currently, surfactant excipients such as polysorbates and poloxamers are used to prevent protein aggregation, which significantly extends shelf-life. Unfortunately, these excipients are themselves unstable, oxidizing rapidly into 100s of distinct compounds, some of which cause severe adverse events in patients. Here, we leverage the highly stable, well-defined, and modular nature of amphiphilic polyacrylamide-derived excipients to isolate the key mechanisms responsible for excipient-mediated protein stabilization. With a library of compositionally identical but structurally distinct amphiphilic copolymers, we quantify multiple relationships between polymer properties and fundamental phenomena to rationally design new ultra-stable surfactant excipients, increasing the stability of a notoriously unstable biopharmaceutical, monomeric insulin, by an order of magnitude. This comprehensive and generalizable understanding of excipient structure-function relationships represents a paradigm shift for the formulation of biopharmaceuticals, moving away from trial-and-error screening approaches towards rational design.

**TOC Graphic:** 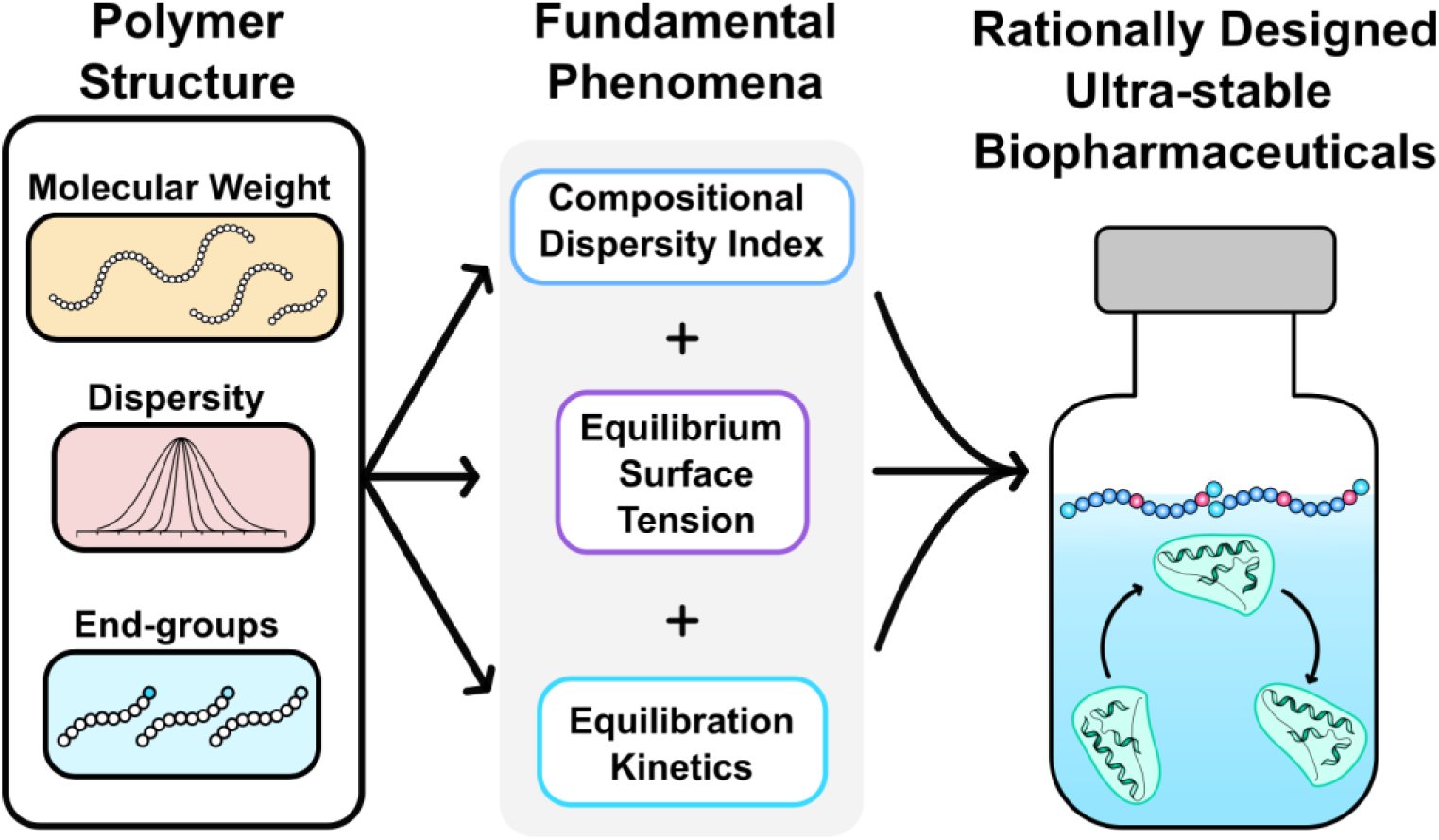

## Introduction

Biopharmaceuticals are the fastest growing class of drugs in the healthcare industry, with the number of products on the market increasing from 13 in 1989 to over 435 in 2022.^1,2^ While the pace of product approvals remained steady until 2015, the rate has tripled from 2017 to 2022, due to the success of antibody and peptide-based therapies increasing the market value from $188 billion to over $343 billion.^2^ The success of these types of biologics is due in large part to complex and delicate tertiary structures that endow these drugs with highly potent and specific biological activity. Unfortunately, these same complex structures make biopharmaceuticals unstable at ambient conditions, necessitating cold chain storage and formulation optimization for clinical translation. Inactive additive technologies, called excipients, have been crucial for the translation of lifesaving drugs from the bench to the bedside.^3^ As proteins are inherently amphiphilic, many are highly prone to aggregation induced by hydrophobic interfaces, such as air or silanized glass, significantly limiting shelf-stability and efficacy.^4,5^ Surfactants, such as polysorbates and poloxamers, are present in >50% of FDA approved biologic drug products and exhibit amphiphilic properties that form protective layers at these hydrophobic interfaces, improving formulation shelf-life.^6^ Recently, these additives have sparked significant safety concerns due to their rapid oxidative degradation and the potential for severe immune reactions, highlighting the critical need to develop alternative next-generation excipients without cleavable esters^7–9^, heterogeneous structures^10–14^, or polyethylene glycol (PEG)^15,16^ functionalities.

Despite this well acknowledged need for new classes of excipients, without a fundamental understanding of how the structural properties of surfactants impact protein stability it remains extremely challenging to design efficacious surfactant excipients *a priori*.^10,17–19^ Specifically, attempts to predict protein stability from structural properties have been thwarted by the highly heterogeneous structure and rapid degradation of current surfactants (pluronics and polysorbates).^11–14,20–22^ Recently, we demonstrated that high-throughput combinatorial synthesis of a large library of amphiphilic polyacrylamide copolymers could enable identification and optimization of a novel class of surfactant excipients capable of dramatically improving the stability of biopharmaceuticals.^4,5,23,24^ Crucially, these polyacrylamide-derived surfactants address the major issues facing current commercial surfactants as they: (i) are structurally well-defined due to their synthesis by reversible addition fragmentation chain transfer (RAFT) polymerization, (ii) are highly stable under diverse conditions (e.g., acidic, basic, oxidative, and enzymatic stress), (iii) do not contain immunogenic PEG, and (iv) exhibit potent surfactant action while not forming micelles.^4,8,9^ By screening this library of novel surfactants with recombinant human insulin stability assays, a specific polymer composition was developed that acted as a potent biopharmaceutical excipient, improving formulation stability and cold-chain resilience over 50-fold compared to standard surfactant excipients.^4,24^ This work was one of the first examples of screening polymer composition to discover new functionality, contributing to the now growing field of high-throughput discovery.^25–32^ These ongoing efforts have seen tremendous success by leveraging the vast chemical space that can be realized by simply combining chemically distinct monomers at different compositional ratios; however, methodologies to understand why these copolymers exhibit improved performance remain sparce. We hypothesized copolymer structural properties such as molecular weight, dispersity of both molecular weight and composition, and end-group functionality, which have historically been neglected in these screening investigations, could provide key mechanistic insights to enable rational design of surfactant excipients. There is a growing appreciation for the importance of these seemingly minor polymer structural characteristics, and indeed end-group functionality has recently been highlighted as a driver of polymer-protein and polymer-interface interactions.^19,33–39^ Moreover, new studies which simulate individual polymer chains highlight that even within systems of compositionally identical polymers, there can be significant differences between structure and function.^32^ With regards to polyacrylamide excipients, there exists a unique opportunity to systematically characterize the impact of molecular weight, dispersity, and end-group functionality on protein stability and to elucidate mechanisms behind these emergent structure-function relationships.

Based on our previous high-throughput screens, we found that a copolymer comprising of acryloylmorpholine (Morph; Mo) and N-isopropylacrylamide (Nipam; Ni) at weight percents of 77% and 23% (MoNi_23%_) was highly effective at stabilizing multiple protein drugs including insulin and antibodies in aqueous formulations under stressed aging conditions. Considering recent efforts to investigate polymer structure-function relationships, we hypothesized that modulating polymer molecular weight, dispersity, and end-group functionality, while controlling for monomer feed composition, would allow us to rationally engineer more stable protein formulations (Figure 1). To evaluate the effect of these properties on excipient performance, we synthesized a library of structurally distinct but compositionally identical polymers and quantified their impact on the stability of monomeric “fast-acting” insulin lispro, a notoriously unstable biopharmaceutical.^4^ We found that molecular weight, dispersity, and end-group functionality each play an important role in the performance of the polyacrylamide excipient and with these structurally distinct copolymers we rigorously investigated of the fundamental phenomena responsible for excipient performance. We discovered that a combination of three functional properties: (i) equilibrium surface tension, (ii) equilibration surface-binding kinetics, and (iii) a new polymer property we call the compositional dispersity index (CDI), accurately explain and predict the enhanced performance of surfactant excipients. Furthermore, leveraging these simple design principles we rationally designed new copolymers – poly(acrylyoylmorpholine-*co-*N-dodecylacrylamide (MoDD) – capable of further improving the stability of monomeric insulin formulations by almost 10-fold compared to MoNi_23%_ and 200-fold compared with polysorbate 80 (PS80), the most common surfactant used in FDA-approved biopharmaceutical drug products (>100h with constant agitation at 37°C). These results are remarkable given that even groundbreaking protein stabilization approaches, such as physical encapsulation, have been shown to stabilize insulin for 6h at room temperature with agitation.^40^ We believe this work generates generalizable design criteria for the development of next-generation excipients by providing quantitative and mechanistic insights into structure-function relationships that dramatically improve the stability of biopharmaceuticals.

**Figure 1.**
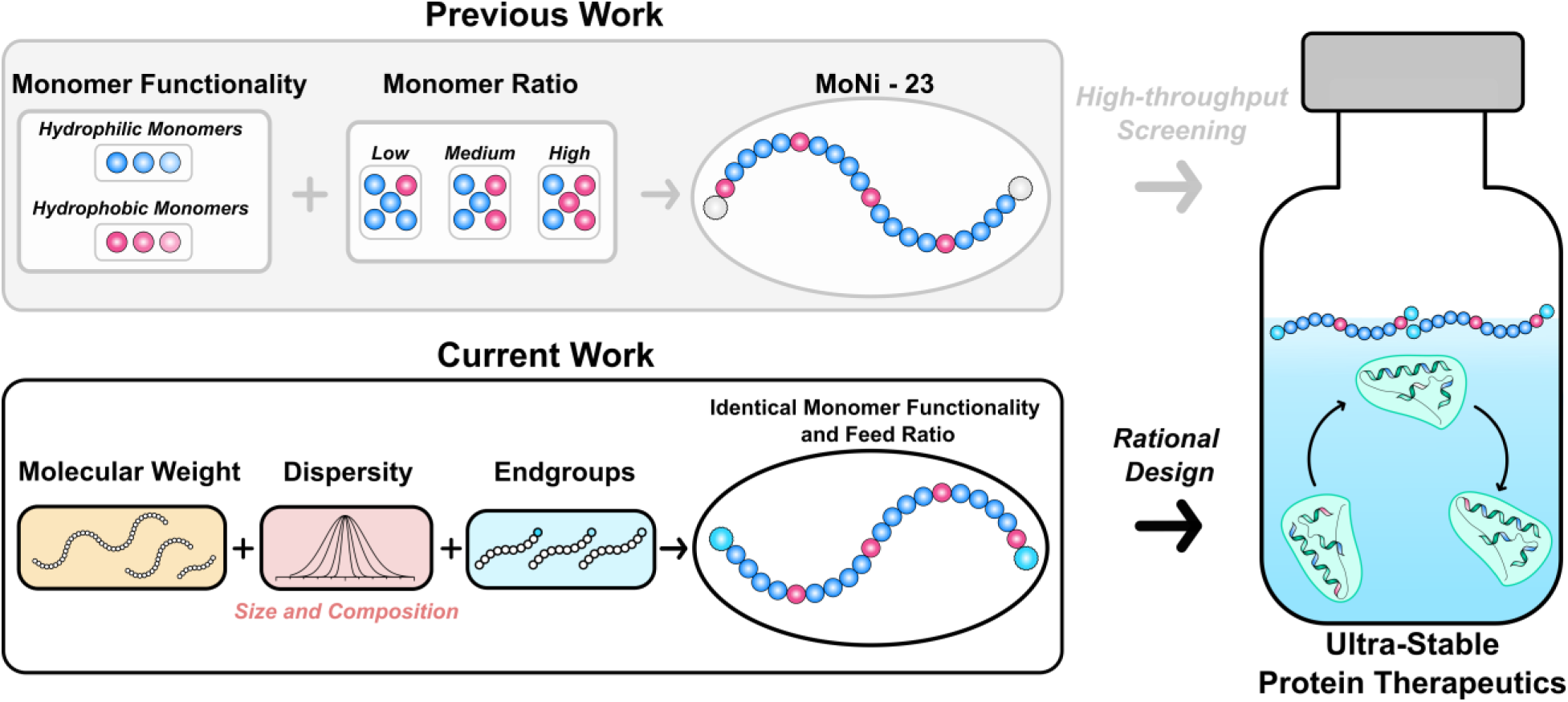
Summary of previous and current polymer property investigations to improve the stability of monomeric insulin formulations.

## Results and Discussion

### Synthesis of MoNi_23%_ analogs

We previously established that a MoNi_23%_ copolymer with a molecular weight of approximately 7 kDa effectively partitions to the air-water interface, precludes protein adsorption at these interfaces, and thereby stabilizes the proteins against aggregation.^4,5,23,24^ To understand the impact of polymer molecular weight, we prepared several copolymers with Mo and Ni incorporation at the previously optimized MoNi_23%_ composition but doped in increasing amounts of chain transfer agent (CTA) (Figure 2A). For these polymerizations we used 2-cyano-2-propyl dodecyl trithiocarbonate (CPDT) as the CTA, which results in low dispersity polymers (Ð < 1.2) even at long reaction times (> 6 h) or low molecular weights (Table 1). Due to the high fidelity of RAFT polymerization, we were able to synthesize six MoNi_23%_ derivatives with identical monomer compositions (Figure 2B), but molecular weights spanning an order of magnitude (2 kDa to 20 kDa) according to size exclusion chromatography (SEC) (Figure 2C). We limited the molecular weights we evaluated to 20 kDa because neutral polymers with molecular weights greater than 25 kDa are too large to be excreted renally by kidney filtration.^41^ While there is significant overlap of the monomer functionalities and the backbone signals in ^1^H NMR, we were able to confirm that all reactions proceeded to high conversion and did not have quantifiable differences in chemical composition (Table 1). Lastly, prior to any stability studies, the CTA end-group was removed via an end-group removal reaction with excess azobisisobutyronitrile (AIBN) and lauroyl peroxide (LPO).^4,5,23,24^

**Figure 2.**
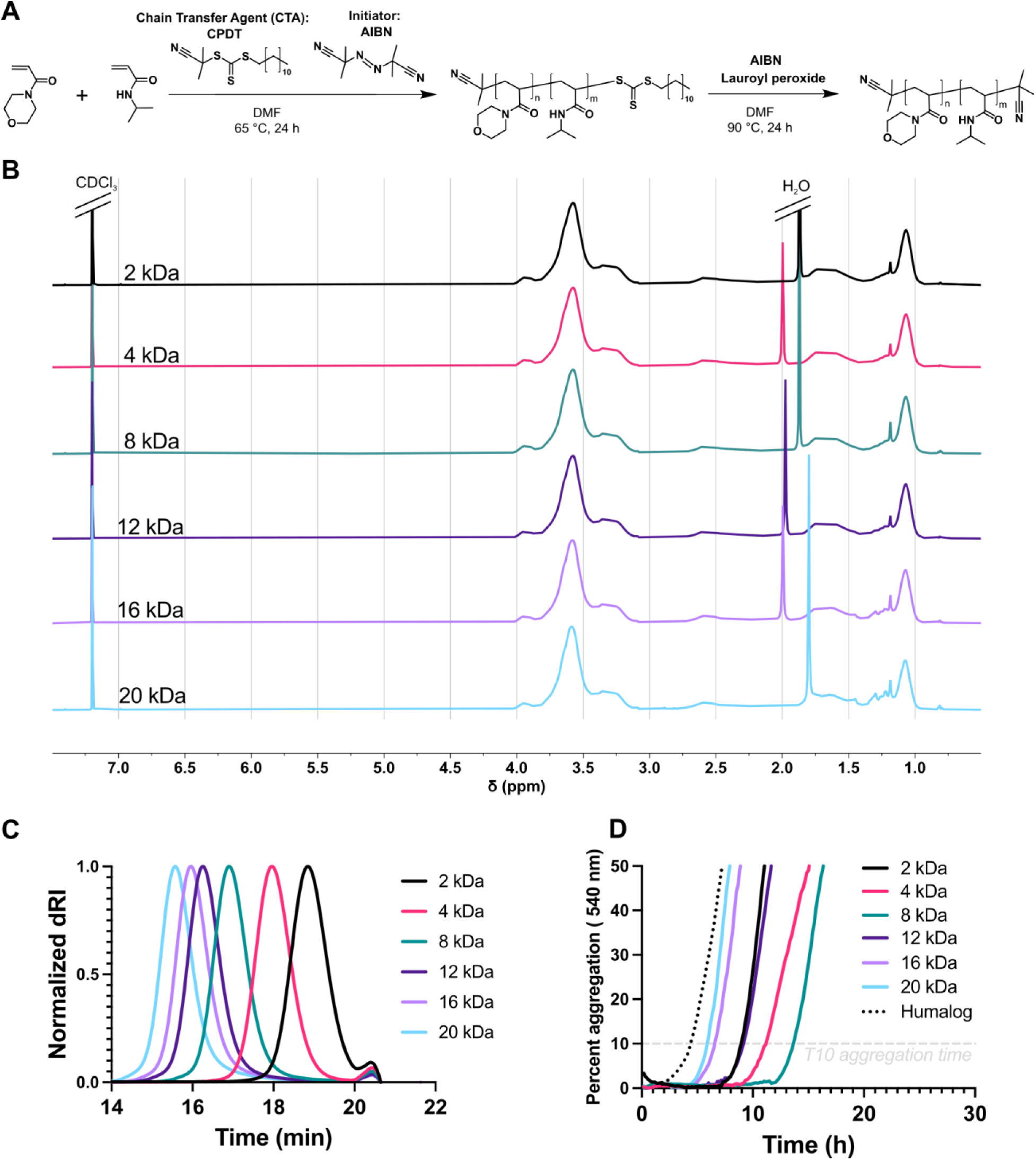
Synthesis, characterization, and evaluation of insulin stability with poly(acrylyoylmorpholine-*co-*N-isopropylacrylamide (MoNi_23%_) excipients at various molecular weights. A) Synthesis scheme for MoNi_23%_ including RAFT polymerization and end-group removal. B) ^1^H NMR characterization of purified MoNi_23%_ polymers of various molecular weights in CDCl_3_. C) Size exclusion chromatography (SEC) determination of MoNi_23%_ molecular weight and dispersity relative to PMMA standards in DMF with LiBr. D) Non-specific absorbance assay to assess stability under stressed aging conditions (37 °C and shaking) of monomeric insulin with EDTA (1.2 mM) + polymer excipients (0.02 wt %) and commercial insulin hexamer (Humalog). Data presented are a mean of 4 technical replicates.

**Table 1.** Summary of MoNi_23%_ characteristics as a function of molecular weight. Counts of Mo and Ni determined from conversion. Conversion* and M_n_* determined with ^1^H NMR. M_n_ and M_w_ determined via SEC with polymethylmethacrylate (PMMA) standards.

| Label | Mo | Ni | Conversion | $M_n^*$ | $M_n$ | $M_w$ | Dispersity |
| --- | --- | --- | --- | --- | --- | --- | --- |
| 2 | 11 | 4 | 98% | 2,400 | 2,000 | 2,100 | 1.08 |
| 4 | 23 | 8 | 97% | 4,500 | 3,600 | 4,000 | 1.11 |
| 8 | 46 | 17 | 97% | 8,700 | 7,600 | 8,400 | 1.11 |
| 12 | 69 | 25 | 98% | 12,900 | 11,600 | 13,300 | 1.14 |
| 16 | 92 | 34 | 96% | 17,100 | 13,700 | 16,200 | 1.19 |
| 20 | 115 | 42 | 97% | 21,300 | 17,500 | 21,200 | 1.20 |
| 8-HD | 33 | 12 | 98% | 6,000 | 6,100 | 9,800 | 1.62 |

### Impact of Molecular Weight

To study the effect of these MoNi_23%_ analogs on protein stability, we made slight modifications to previously published assays to increase throughput and accentuate the differences between polymers.^4^ In brief, we used the zinc chelating agent, ethylenediaminetetracetic acid (EDTA), at a concentration of 1.2 mM to convert hexameric commercial insulin into insulin monomers and added polymeric excipients at 0.2 mg/mL (0.02 wt%) before heating to 37 °C and shaking continuously. Insulin aggregation was measured by monitoring non-specific absorbance at 540 nm with the time to 10% of maximum absorbance being used as the onset of aggregation (T10 aggregation time).^42^ All polymers stabilized monomeric insulin formulations for longer than the commercially available hexameric insulin formulation, trade name Humalog (Figure 2D). Surprisingly, we found that intermediate molecular weights (4 kDa and 8 kDa) stabilized the biopharmaceutical longer than both the low (2 kDa) and high molecular weight (12 kDa, 16 kDa, and 20 kDa) polymers (Figure 2D). While this result explains the success of the original MoNi_23%_ (7 kDa), it was unexpected because literature precedent implies that surface properties of polymeric surfactants vary linearly with molecular weight, either improving at low molecular weights due to increased packing efficiency or at higher molecular weights due to avidity effects.^38,43^ This unique phenomenon of an optimal insulin stability at an intermediate molecular weight hinted at the possibility that our current understanding of excipient-mediated stabilization of proteins is incomplete, and that several material properties may be important for this application.

### Impact of Molecular Weight Dispersity

As a secondary method to validate the observed effect of molecular weight on insulin stability, we varied polymer dispersity at a fixed molecular weight. If the most “active” components of the polyacrylamide-derived surfactant are the polymer chains with the moderate molecular weights (in the range of 4-8 kDa), we hypothesized that at higher dispersities, where these weight fractions would be lower total percentages of the polymer, the insulin stability would decrease. To synthesize high dispersity (HD) polyacrylamides, Anastasaki and coworkers have demonstrated that 2-Cyanopropan-2-yl *N*-methyl-*N*(pyridin-4-yl)carbamodithioate (CMCD) could be used to achieve a target molecular weight with high chain-end retention.^44,45^ We replicated these protocols to synthesize compositionally identical 4 kDa and 8 kDa MoNi_23%_ analogs with dispersities of 1.33 and 1.62 respectively (Figure 3A and S1A). The end-groups of the HD polymers were removed via several radical reactions to ensure complete removal of the CTA (Table 2 and Table S1, Figure S2-5) and mixed with the low dispersity counterparts at various ratios to create new polymers with similar number average molecular weights but varied dispersities (Figure 3B and Table 2). We observed consistent, quantifiable, and incremental decreases in the T10 aggregation time of monomeric insulin as a function of MoNi_23%_ dispersity (Figure 3C), which was consistent for both 8 kDa and 4 kDa polymers (Figure S1B). These results highlight that while polymers may appear identical at the bulk level (composition and molecular weight) differences at the level of the individual chains in the population (dispersity) can result in distinct functional behaviors. This report is the first to systematically quantify the effect of polymer dispersity in the context of biomaterial optimization and demonstrate its importance as a tunable property.

**Figure 3.**
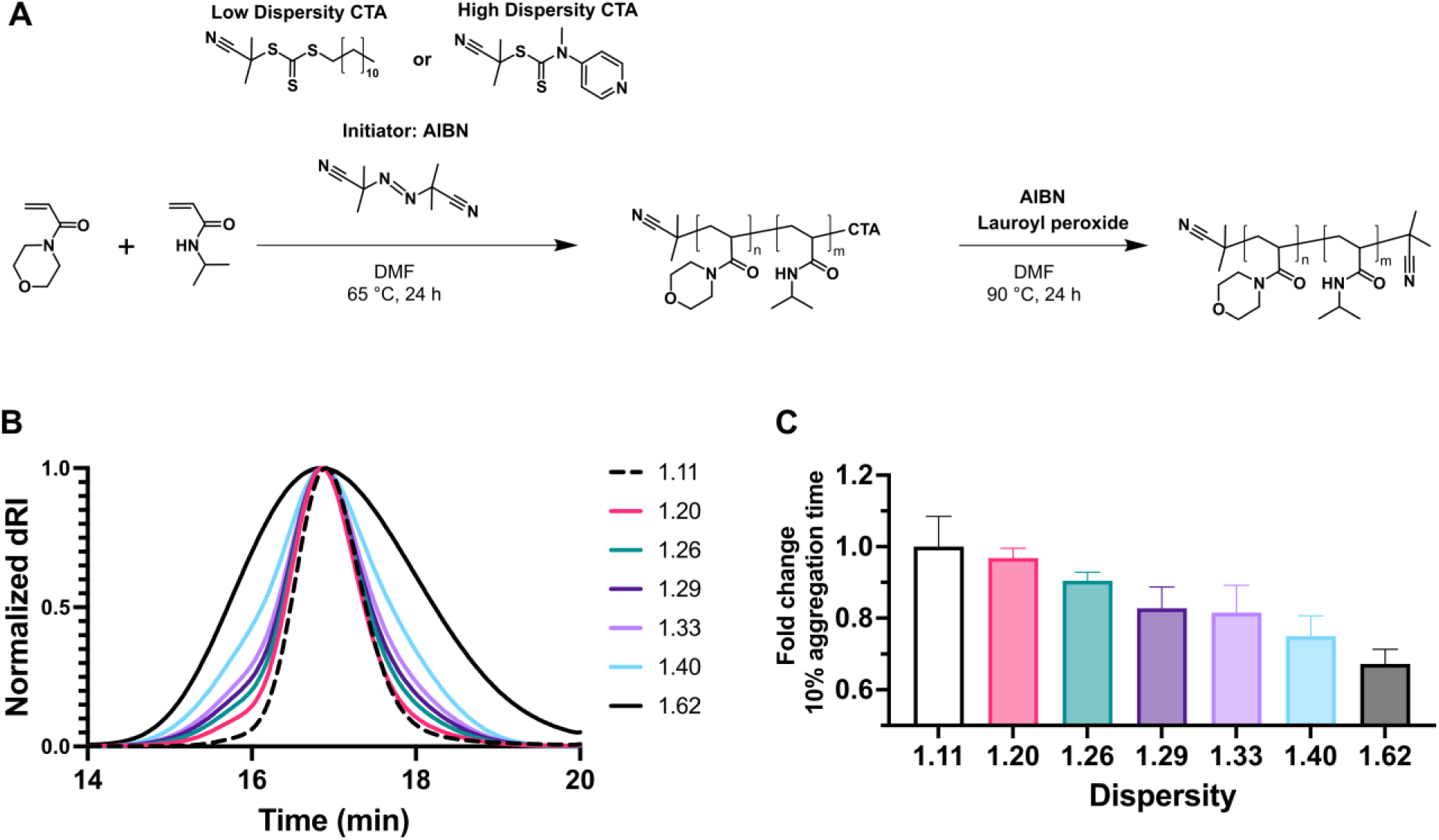
Synthesis, characterization, and evaluation of insulin stability with MoNi_23%_ excipients at various dispersities. A) Synthesis scheme for 8 kDa MoNi_23%_ polymers with identical compositions but different dispersities. B) SEC traces for mixtures of high and low dispersity MoNi_23%_ polymers yielding a gradient of dispersities. C) Comparative T10 insulin aggregation times relative to low dispersity (Đ=1.11) 8 kDa MoNi_23%_ as a function of dispersity. Data presented is a mean of 4 technical replicates.

**Table 2.** Evaluation of dispersity of MoNi_23%_ 8 kDa and 8-HD mixtures with SEC and PMMA standards. End-group removal was determined by ratio of the area under the curve (AUC) before and after the reaction at 310 nm.

| Label | M<sub>n</sub> | M<sub>w</sub> | Dispersity | End group removal |
| --- | --- | --- | --- | --- |
| 8 | 7,575 | 8,433 | 1.11 | 98.3% |
| 1.20 | 7,639 | 9,127 | 1.20 | --- |
| 1.26 | 7,441 | 9,345 | 1.26 | --- |
| 1.29 | 7,357 | 9,499 | 1.29 | --- |
| 1.33 | 7,226 | 9,607 | 1.33 | --- |
| 1.40 | 7,071 | 9,919 | 1.40 | --- |
| 8-HD | 6,072 | 9,815 | 1.62 | 98.47% |

### Impact of End Groups

To investigate the effect of excipient chain ends on insulin stability, we chose to first study the best performing low molecular weight polymer (MoNi_23%_ 4 kDa) since we believed the greater proportion of end-groups would magnify the differences between various chemical functionalities. For all MoNi_23%_ analogs synthesized to this point, the end group removal reaction was performed with AIBN, in accordance with previously established protocols.^4,5,23,24^ We continued to leverage the high efficiency of excess radical end-group removal reactions to synthesize new MoNi_23%_ analogs with 2,2’-Azobis(4-methoxy-2,4-dimethylvaleronitrile) (V-70), 2,2’-Azobis[2-(2-imidazolin-2-yl)propane]dihydrochloride (VA-044), and 4,4′-Azobis(4-cyanopentanoic acid) (ACVA) and compare their activity to 4 KDa MoNi_23%_ with AIBN. These initiators cover a range of chemical functionalities at physiologic pH, with V-70 being more hydrophobic than AIBN, ACVA adopting a negative charge, and VA-044 adopting a positive charge (Figure 4A). While all end-group removal reactions achieved greater than 95% efficiency as quantified via absorbance at 310 nm (Table 3 and Figure S6), the chemical identity of the thermal initiator had a significant impact on the performance of the polymeric excipient.

**Figure 4.**
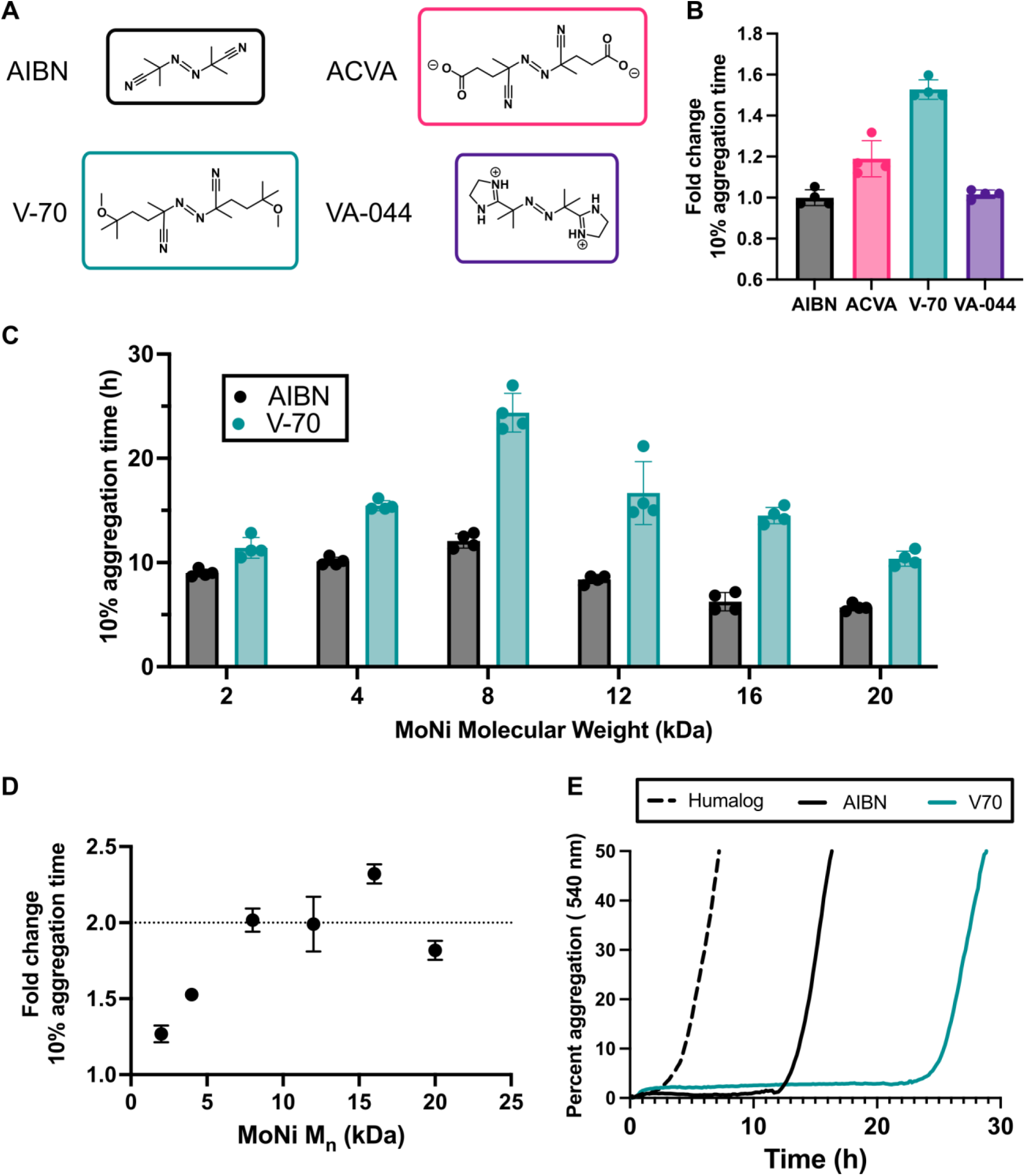
Comparison of various thermal initiators that were used for end-group removal and their impact on insulin stability. A) Structures of thermal initiators and their ionization states in insulin formulations. B) Comparative T10 insulin aggregation times relative to 4 kDa MoNi_23%_ with AIBN end-groups. C) T10 insulin aggregation times compared at various molecular weights with either AIBN or V-70 end-groups. D) Ratio of T10 insulin aggregation times of V-70 to AIBN end-groups as a function of MoNi_23%_ molecular weight. E) Representative non-specific absorbance assay of monomeric insulin stabilized with 8 kDa-AIBN and 8 kDa-V70 vs. commercial Humalog. Data presented are a mean of 4 technical replicates.

**Table 3.** Summary of end-group removal efficiency of MoNi_23%_ 4 kDa with various initiators as determined by AUC at 310 nm during SEC.

| Label | End group removal |
| --- | --- |
| 4-AIBN | 99.2% |
| 4-ACVA | 99.1% |
| 4-V70 | 98.9% |
| 4-VA | 98.0% |
| 8-AIBN | 98.3% |
| 8-V70 | 95.9 % |

Specifically, the neutral initiator V-70 dramatically increased the T10 aggregation time of the 4 kDa MoNi_23%_ by more than 1.5-fold (Figure 4B and S7). To investigate possible mechanisms and understand the impact of end-group identity as a function of molecular weight, all six polymers from previous experiments were functionalized with V-70. Contrary to our hypothesis, we observed that the switch from AIBN to V-70 increased the T10 aggregation time of the high molecular weight polymers by 2-fold, proportionally more than their low molecular weight counterparts, which only saw increases of ∼1.5-fold (Figure 4C, 4D, and S10). To underscore the impact of simply modulating the end-capping reagent, we show that by SEC and NMR the 8 kDa MoNi_23%_ with AIBN and V-70 end-groups are essentially identical (Figure S8 and S9) yet the 8 kDa MoNi_23%_ with V-70 stabilizes insulin for over 24 h compared to the 8 kDa MoNi_23%_ with AIBN, which aggregates in 12 h (Figure 4E). These results demonstrate that regardless of molecular weight, the chain-end identity of surfactants plays a vital role in their performance as stabilizing excipients.

### Mechanistic Investigations of Surface Activity

We previously discovered that for both low and high concentration biopharmaceuticals, MoNi_23%_ significantly reduced the surface tension of the formulation. These results suggested that the mechanism behind the increased protein stability was a preferential accumulation of polymer at the air-water interface preventing the proteins from denaturing upon interacting with the hydrophobic air.^4,23,24^ We suspected that the observed differences in insulin stability as a function of molecular weight, dispersity and end-groups would correspond with lower surface tension values. We chose to evaluate the surface tensions of our best performing polymers (low dispersity with V-70 end-groups) at various molecular weights with the Wilhelmy Plate method (Figure 5A). We observed a monotonic trend in surface tension as a function of the polymer molecular weight with the 2 kDa MoNi_23%_ equilibrating at the lowest surface tension of 44.88 mN/m and the 20 kDa MoNi_23%_ equilibrating at the highest surface tension of 52.29 mN/m (Figure 5B). Combined with our previous insulin stability experiments (Figure 4C) demonstrating that the 2 kDa and 20 kDa MoNi_23%_ polymers have the lowest T10 aggregation time, it becomes clear that surface tension alone is insufficient to explain the performance of polymeric excipients as stabilizers of biopharmaceuticals. Moreover, despite the more than 2-fold difference in T10 aggregation time between 8 kDa MoNi_23%_ with V-70 and AIBN end-groups (Figure 4E), the equilibrium surface tension of these polymers is almost identical at 49.88 mN/m and 50.92 mN/m (Figure 5C and Table 4). When we plotted the T10 aggregation time vs. surface tension (Figure 5D), no correlation emerged, corroborating our observations that surface tension alone cannot account for increases in protein stability.

**Figure 5.**
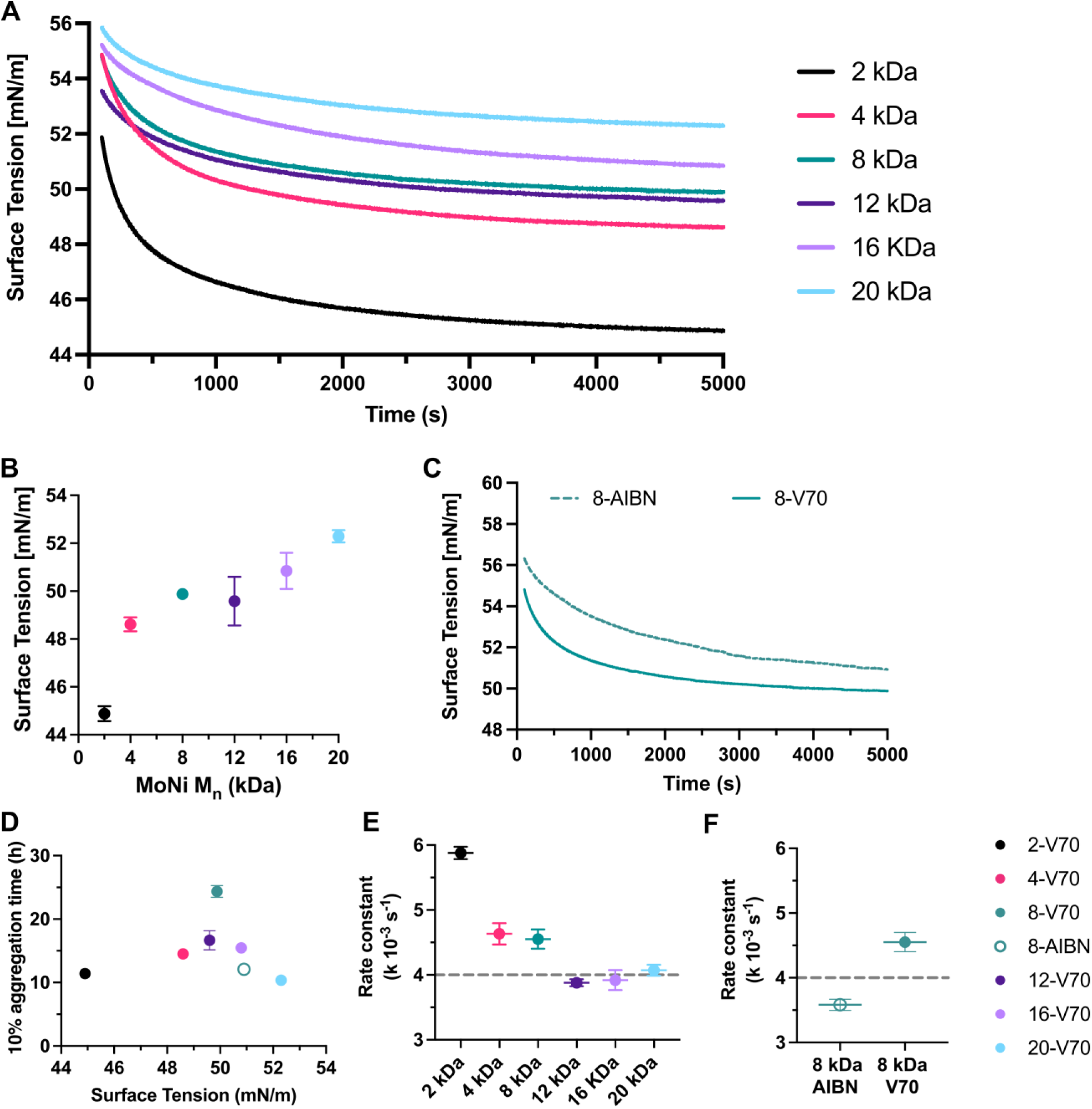
Evaluation of surface tension with Wilhelmy plate method as a function of MoNi_23%_ molecular weight and end-group identity. A) Mean surface tension values of 0.02 wt % MoNi_23%_ in Humalog buffer (without insulin) over 5000 seconds (s) (n = 2). B) Equilibrium surface tension of solutions with MoNi_23%_ excipients (0.02 wt %) at 5000 s as a function of polymer molecular weight. C) Surface tension over 5000 s of 8 kDa MoNi_23%_ with either AIBN end-group or V-70 end-group. D) T10 aggregation times for insulin stabilized with different MoNi_23%_ molecular weights as a function of equilibrium surface tension. E) Rate constant for a one phase exponential decay fit of the surface tension equilibration measurements in the first 300 s as a function of MoNi_23%_ molecular weight. F) Rate constant for a one phase exponential decay fit of the surface tension equilibration measurements in the first 300 s for MoNi_23%_ with either AIBN end-group or V-70 end-group. Mean and standard error of the mean were obtained from duplicate experiments.

**Table 4.** Equilibrium surface tension means and standard errors for duplicate measurements of 0.02 wt % solutions of MoNi_23%_ polymers in Humalog Buffer as measured by the Wilhelmy plate method. Rate constants from one phase exponential decay of surface tension measurements in the first 5 min of equilibration. Compositional Dispersity Index (CDI) for each polymer determined from Compositional Drift simulations. Summary of T10 aggregation times for each polymer evaluated by insulin stability assay.

| Label | Equilibrium Surface Tension (mN/m) | Surface Kinetics Rate constant ($k \cdot 10^{-3} \text{ s}^{-1}$) | Compositional Dispersity Index | 10% Aggregation Time (h) |
| --- | --- | --- | --- | --- |
| 2-V70 | 44.9 ± 0.3 | 5.878 ± 0.047 | 7.86 | 11.42 ± 0.50 |
| 4-V70 | 48.6 ± 0.2 | 4.633 ± 0.082 | 5.54 | 15.46 ± 0.24 |
| 8-V70 | 49.88 ± 0.02 | 4.551 ± 0.074 | 2.10 | 24.37 ± 0.93 |
| 12-V70 | 49.6 ± 0.7 | 3.880 ± 0.027 | 1.56 | 16.67 ± 1.51 |
| 16-V70 | 50.8 ± 0.5 | 3.919 ± 0.077 | 1.25 | 14.5 ± 0.39 |
| 20-V70 | 52.3 ± 0.2 | 4.070 ± 0.042 | 1.20 | 10.37 ± 0.36 |
| 8-AIBN | 50.9 ± 0.6 | 3.583 ± 0.044 | 2.10 | 12.08 ± 0.35 |
| Humalog Buffer | 71.960 ± 0.002 | N/A | N/A | N/A |

While equilibrium surface tension did not yield clear correlations with T10 aggregation times, we noticed differences in the equilibration kinetics during these time-resolved experiments, particularly at short time points (< 5 min, ∼300s). We fitted our surface tension curves with a one phase exponential decay curve and extracted rate constants (Table 4 and Figure S11) to quantify how quickly these surfactants partition to the air-water interface. These rate constants roughly decreased as a function of molecular weight (Figure 5E), with larger polymers achieving equilibrium surface tensions at slower rates than smaller polymers. Similar to the equilibrium surface tension measurements, there was not a simple correlation with protein stability and these rate constants since the smallest 2 kDa MoNi_23%_ which is one of the worst performing excipients, had the fastest rate. Yet, when we compared the rate constants for 8 kDa MoNi_23%_ polymers with either AIBN or V-70 end-groups, the V-70 polymers equilibrated much faster than their AIBN counterparts (Figure 5F and S12). This observation implies that above a certain molecular weight threshold (above 4 kDa for MoNi_23%_), the faster the surfactant partitions to the air-water interface the greater the enhancement of insulin stability. Importantly, this observation explains why higher molecular weight polymers with slower kinetics benefit more from end group optimization (Figure 4D), as low molecular weight polymers can already achieve equilibrium rapidly due to their small size there is nothing to be gained from the end group substitution.

### Mechanistic Investigation of Compositional Dispersity

We were struck by the existence of a clear low molecular weight threshold for these polyacrylamide-derived excipients, below which, despite excellent surface-active properties, the excipients lost the ability to effectively stabilize insulin. While our polymerization reactions contained identical monomer stoichiometry, the stochastic nature of copolymerizations can lead to a range of possible polymer compositions for each individual polymer chain. Monte Carlo simulations have emerged as a powerful tool to understand the composition of individual chains within a polymerization.^32,46,47^ In particular, we noticed that at low molecular weights individual polymer chains deviate substantially from the stoichiometrically ideal copolymer composition.^48^ To explore this characteristic among the MoNi_23%_ variants in our study, monomer ratios, reactivity ratios, and dispersities were used to simulate each polymerization at a given molecular weight with a previously published Monte Carlo simulation package, Compositional Drift (Figures S13-18). We plotted the total degree of polymerization (DP) of each monomer within a single polymer chain to create novel 2D visualizations of the overall compositional dispersity (Figure 6A). An ideal polymer chain at any given molecular weight would have a ratio of approximately 3 Morph monomers to 1 Nipam monomer, creating a line from the origin with a slope of about 1 over 3 (0.299 to be exact). Therefore, the compositional dispersity of the simulated polymerization can be assessed by how well the distribution of individual polymer chains fits this ideal copolymer line, quantified by R^2^. This approach enabled us to quantify the compositional dispersity of each polymer and we found that dispersity decreased at higher molecular weights (higher R^2^ values) indicating that more polymers in the simulation achieved a composition close to the ideal.^48^ In fact, we observed a similar molecular weight cut-off as in our insulin stability study, due to the dispersity becoming significantly higher below 4kDa. We chose to describe these dispersity values on a scale analogous to molecular weight as compositional dispersity index (CDI, |1/R^2^|), yielding values of 7.86, 5.54, 2.1, 1.56, 1.25, 1.20 for 2 kDa, 4 kDa, 8 kDa, 12 kDa, 16 kDa and 20 kDa, respectively (Figure 6B). This large spread of individual polymer compositions explains the poor ability of low molecular weight polyacrylamide-based excipients to stabilize insulin, as our previous work demonstrated that a composition of exactly 77% Morph and 23% Nipam was key to excipient performance.^4^ Moreover, at high compositional dispersity, extreme outliers in composition might seed insulin aggregation by inducing amyloid fibril formation in a mechanism similar to short amphiphilic peptides.^49–51^ In addition to explaining the inability of short MoNi_23%_ polymers to effectively prevent protein aggregation, compositional dispersity also explains the slight differences in the performance of 4 kDa and 8 kDa MoNi_23%_ polymers. Despite almost identical surface properties (Table 4), 8 kDa MoNi_23%_ stabilizes insulin for 1.5 times longer than 4 kDa MoNi_23%_. Only when we compare the CDIs, 2.1 vs. 5.54, does it become clear why the 8 kDa MoNi_23%_ polymer so significantly outperforms its 4 kDa counterpart. In summary, we provide a new metric, CDI, quantifying heterogeneity in the distribution of individual polymer compositions to elucidate subtle differences between polymerizations with identical feed ratios. This metric provides a key insight into the performance of excipients and when combined with measurements of surface properties can explain how a surfactants structure ultimate impacts protein stability.

**Figure 6.**
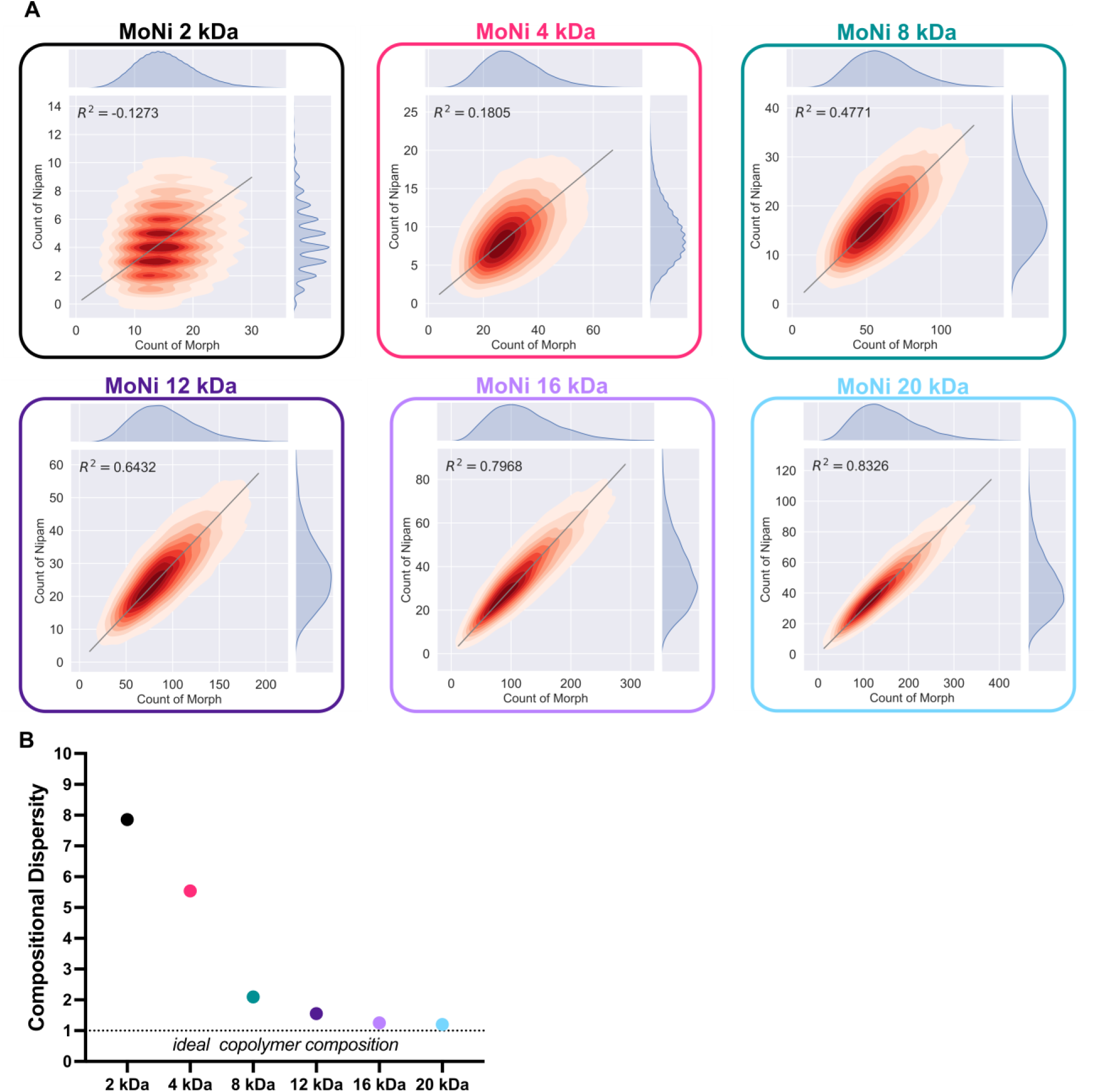
Compositional dispersity plots generated from Compositional Drift simulation. A) 2D-plots of incorporation of Morph and NIPAM simulated for individual polymer chains to visualize monomer compositions of MoNi_23%_ polymers with various molecular weights. To quantify deviation from the ideal monomer composition based on feed ratio (Morph/NIPAM of 77/23), R^2^ was calculated for the distributions against a line with a slope of 23/77. B) Compositional Dispersity Index (defined as |1/R^2^|) as a function of MoNi_23%_ molecular weight.

### Fundamental Design Principles for Effective Excipients

Through our mechanistic investigations, we determined that surface tension, equilibration kinetics, and compositional dispersity index (CDI) can explain the performance of copolymer excipients. With these factors in mind, we propose a framework for designing copolymer excipients that maximizes protein stability (Figure 7A). First, copolymers must be of sufficient molecular weight to have a homogenous monomer composition at the individual polymer level (low CDI). Second, copolymers must be small enough to pack tightly at the air-water interface (as determined by equilibrium surface tension). Third, given copolymers with similar compositional dispersities and equilibrium surface tensions, the highest rate constant for surface equilibration will provide the most protection against protein aggregation. Using the MoNi_23%_ library as a case study, we can begin to understand trends in insulin stability as a function of each of these mechanisms (Figure 7B). When we plot CDI vs. a surface property (such as rate constant), clusters with high and low T10 aggregation times emerge. However, with only two properties it is still difficult to account for all the variation observed between polymers. Applying the above framework, we can isolate a range of molecular weight values over which each mechanism plays the dominant role in predicting insulin stability (Figure 7C). At low molecular weights (at and below 8 kDa) CDI explains 99% of the variation in the T10 aggregation time (Figure 7D). At high molecular weights (at or above 12 kDa), equilibrium surface tension explains 99% of the variation in insulin stability (Figure 7E). Finally, at intermediate molecular weights (between 8 kDa and 16 kDa) the rate constant for equilibration explains 96% of the variation in T10 aggregation times, including differences between end-groups (AIBN vs. V70). The ability of this rate constant to predict insulin stability in this group of extremely similar polymers (8 kDa-V70, 12 kDa-V70, 16 kDa-V70, and 8 kDa-AIBN), which all had compositional dispersities below 3 and equilibrium surface tensions values between 49.9 and 50.9 mN/m, is strong evidence that we have developed a rigorous understanding of the fundamental principles underlying polymeric excipient performance. Taken together, we have clearly defined three regimes where CDI, equilibrium surface tension, and equilibration kinetics can explain and predict the performance of copolymer excipients.

**Figure 7.**
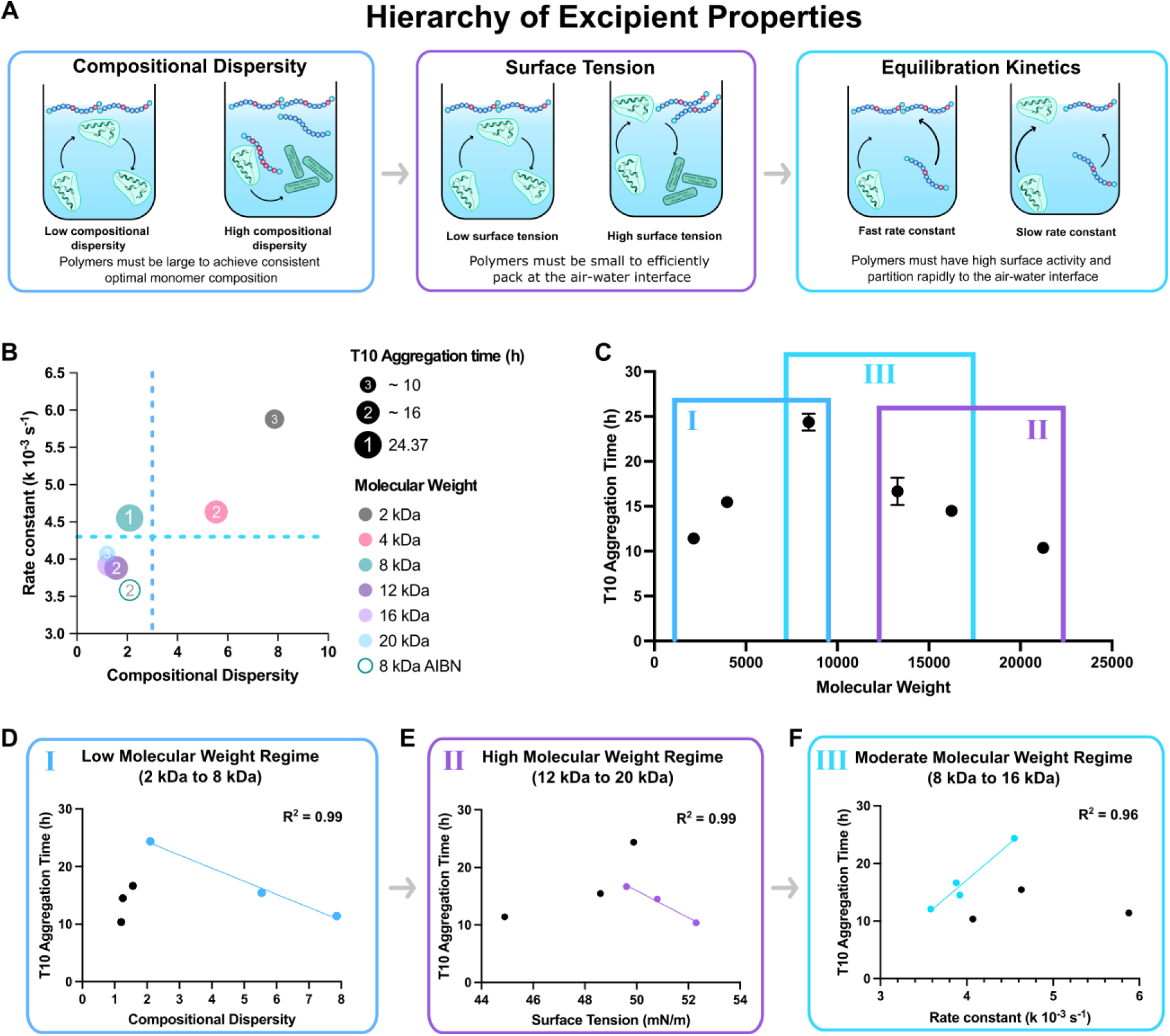
A) Schematic overview summarizing excipient properties and mechanistic explanation of their impact on protein stability. Excipients must first have low compositional dispersity, then low surface tensions, and finally fast equilibration kinetics to prevent aggregation of monomeric insulin. B) Comparison of rate constants, compositional dispersity, and T10 aggregation time for various MoNi_23%_ polymers. C) Selection of 3 molecular weight regimes and the key polymer property that explains and predicts excipient performance (T10 aggregation time). D) Linear correlation between Compositional Dispersity Index and T10 aggregation time at low molecular weights. E) Linear correlation between Surface Tension and T10 aggregation time at high molecular weights. F) Linear correlation between rate constant and T10 aggregation time for polymers with low compositional dispersity and low surface tensions (moderate molecular weights). Fits performed with Prism 10.

### Rationale Design of Next-Generation Surfactant Excipients

With the well-defined and modular characteristics of our copolymer library we were able to quantitatively define structure-function relationships and ultimately predict excipient performance with a rigor that was previously not feasible due to the heterogeneity and rapidly degrading nature of commercially available excipients. To demonstrate the utility of our hierarchy of excipient properties, we designed and synthesized a series of chemically distinct surfactant excipients with high surface activity and low CDI, enabling ultra-stable monomeric insulin formulations (T10 exceeding 100 h). Since homogeneity and surface activity (Figure 7B) are the key properties of high performing excipients, we designed high molecular weight (10 kDa or greater) copolymers with a new hydrophobic monomer, dodecylacrylamide (Dodecyl; DD). This monomer contains a single intermediate-length carbon chain, which is a standard surfactant tail for many commercial excipients (e.g., polysorbate 20) and was recently established as one of the most effective tail moieties for antibody stabilization with surfactants.^19^ Additionally, we opted to not remove the end-groups of these polymers in an effort to demonstrate that surface activity can be controlled by both end-group identity and copolymer composition (Figure 8A). Since the hydrophobicity of the dodecyl monomer is significantly higher than the Nipam in MoNi_23%_ and most commercial excipients only require a single surfactant tail, we chose to incorporate 5 DD monomers per polymer to balance solubility and surface activity. Prior to synthesis, we simulated the compositional dispersity of copolymers comprising acryloylmorpholine (Morph; Mo) and dodecylacrylamide (Dodecyl; DD) and easily found two copolymers at 10 kDa (MoDD10) and 20 kDa (MoDD20) exhibiting CDIs below 3 (Figure 8C, S19, S20, and S21). It is noteworthy that MoDD20 had a higher CDI (2.02) than the MoDD10 (1.64), despite having a higher molecular weight, because of the lower weight percent loading of the dodecylacrylamide monomer (5% vs. 10%). These new simulations highlight that CDI is a function of both degree of polymerization and comonomer feed ratio, increasing whenever the DP of one monomer approaches low single digit values.

**Figure 8.**
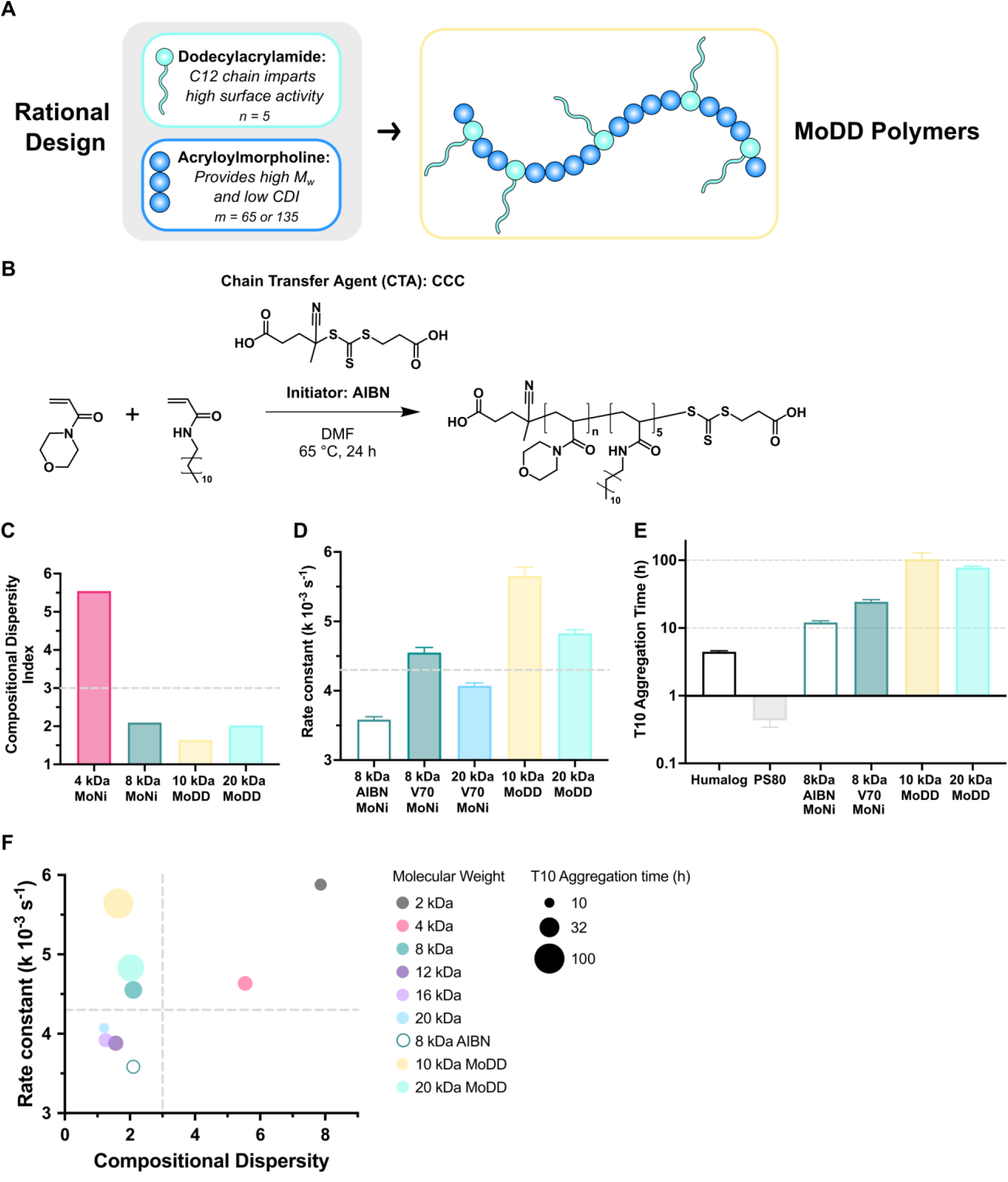
A) Simple rational design of new excipients based on hierarchy of excipient properties. B) Synthesis of poly(acryloylmorpholine-*co-*N-dodecylacrylamide (MoDD)polymers. C) Compositional Dispersity Index for MoDD polymers compared to MoNi_23%_analogs via Monte Carlo simulations with Compositional Drift. D) Rate constant for a one phase exponential decay fit of the surface tension equilibration measurements in the first 300 s for MoDD polymers compared to MoNi_23%_analogs. Mean and standard error of the mean were obtained from duplicate experiments. E) T10 monomeric insulin aggregation times for MoDD polymers compared to commercial products (Humalog and PS80) and MoNi_23%_ analogs. Data presented are mean and standard error. F) Comparison of rate constants, compositional dispersity, and T10 aggregation time for various MoDD and MoNi_23%_ polymers. Copolymers with high surface engagement rate constants and low compositional dispersity (top left quadrant) exhibit the highest T10 aggregation times.

With these validated compositions in hand, we synthesized the two MoDD polymers analogously to the described RAFT polymerizations for MoNi_23%_ (Figure 8B). The polymerizations ran to high conversion (98%) and the copolymers exhibited monomodal SEC traces with low molecular weight dispersity (Figure S22), showing only slight tailing in the SEC likely due to the dodecyl moieties interacting with the column. When we evaluated the surface activity (Figure S23) of MoDD10 and MoDD20, both copolymers exceeded the 4.3 x 10^-3^ s^-1^ equilibration rate constant cut off established by our MoNi_23%_ studies (Figure 8D and S24). Furthermore, despite comprising dodecyl moieties, these copolymers did not hydrophobically self-assemble into micelles at concentrations up to 5 mg/mL (Figure S25).

Finally, to assess the performance of these rationally designed excipients we compared the commercial “fast-acting” insulin formulation Humalog, monomeric insulin formulations stabilized with either a common commercial surfactant excipient (PS80), our previous leading MoNi_23%_ derivative (8 kDa AIBN), our optimized MoNi_23%_ (8 kDa V70), or our new MoDD excipients (MoDD10 and MoDD20). In these studies, PS80-stabilized monomeric insulin formulations were less stable that commercial Humalog (Figure 8E and S26). In contrast, our rationally designed excipients MoDD10 and MoDD20 increased the stability of monomeric insulin formulations to over 100 h under these stressed aging conditions (20-fold compared to Humalog). These surfactants improved formulation stability by over 200-fold compared to PS80-stabilized formulations and by roughly 10-fold compared to MoNi_23%_ (Figure 8E and S27, Table S2). When placed in context of our surfactant excipient design criteria (Figure 8F), both MoDD10 and MoDD20 fall solidly within the goldilocks region of low compositional dispersity and rapid surface engagement. Leveraging our mechanistic understanding of excipient performance, we designed and synthesized two chemically distinct next-generation excipients enabling ultra-stable monomeric insulin formulations.

## Conclusion

In this study we establish a comprehensive framework to predict the performance of surfactant excipients and demonstrate that use of these criteria can replace high-throughput screening for the identification of next-generation excipients. We leveraged the modular and well-defined characteristics of polyacrylamide surfactants to isolate and measure new variables for excipient optimization. With quantification of surface activity and the development of a new polymer descriptor, compositional dispersity index (CDI), we defined three molecular weight regimes where the stability of monomeric insulin could be linearly explained by a single dominant excipient property. While large trial-and-error screens of surfactants have been performed previously, systematic investigations of structure-function relationships were not feasible or productive to due to the highly unstable and heterogenous nature of commercial excipients. For the first time for any class of excipients, we created a framework that enables accurate prediction of excipient performance from fundamental principles. From this understanding of fundamental phenomena, we were able to rationally and efficiently engineer the structure and consequently, the activity, of amphiphilic copolymer excipients via several distinct mechanisms: (i) minimizing heterogeneity (dispersity) in both molecular weight and composition increased the T10 aggregation time by approximately 1.5-fold and 2-fold respectively, (ii) molecular weight optimization to reduce equilibrium surface tension more than doubled the stability of monomeric insulin formulations compared to commercially available insulin, Humalog (from 4.5 h to 12 h), and (iii) decreasing equilibration time via end-group optimization again doubled the T10 aggregation time (from 12 h to 24 h). Harnessing these generalizable design criteria, we synthesized two novel surfactant excipients, MoDD10 and MoDD20, and showed they could improve monomeric insulin formulation stability by orders of magnitude compared to current commercial “fast-acting” insulin drug products like Humalog. Importantly, all these stability assays were under extremely harsh stressed aging conditions, implying that under less austere conditions like room temperature (Humalog is labeled as stable for ∼ 30 days), these monomeric insulin formulations may be stable for upwards of a year. Such an improvement would represent a major milestone for the clinical translation of ultra-fast acting monomeric insulin drugs as stability and cold-chain storage are critical hurdles hindering clinical translation. In summary, we utilized the distinct structure-function relationships in a library of amphiphilic copolymers to dramatically increase the stability of therapeutically relevant monomeric insulin and define key, generalizable design criteria for high performance excipients as stabilizers in biopharmaceutical formulations.

## Acknowledgements

This work was supported in part by the JDRF (2-SRA-2022-1168-M-B). A.N.P. was supported by a Stanford Maternal and Child Health Research Institute postdoctoral fellowship. L.T.N. was supported by a Stanford Graduate Fellowship in Science and Engineering. N.E. was supported by a National Science Foundation Graduate Research Fellowship (DGE-2146755). Special thanks to Nicole Fox for consulting on figure design. We would also like to thank members of the Appel lab, especially Julie Baillet, Andrea I. d’Aquino, Shoshana Williams, Christian Williams, and Sophia Bailey for their helpful discussions.

## Author Contributions

Conceptualization, A.N.P. and E.A.A.; Methodology, A.N.P., L.T.N., S.B., G.G.F., and E.A.A.; Data Analysis and Simulations, A.N.P., N.E., S.T., A.A.A.A., and E.A.A.; Investigation, A.N.P.; Writing, A.N.P., and E.A.A.; Editing, A.N.P., N.E., L.T.N., S.B., A.A.A.A., and E.A.A. Visualization A.N.P., N.E., L.T.N., S.T., and E.A.A.; project administration, E.A.A.; funding acquisition, A.N.P. and E.A.A.

## References

(1) Mitragotri, S.; Burke, P. A.; Langer, R. Overcoming the Challenges in Administering Biopharmaceuticals: Formulation and Delivery Strategies. Nature Reviews Drug Discovery 2014, 13 (9), 655–672. 10.1038/nrd4363.

(2) Walsh, G.; Walsh, E. Biopharmaceutical Benchmarks 2022. Nat Biotechnol 2022, 40 (12), 1722–1760. 10.1038/s41587-022-01582-x.

(3) Vargason, A. M.; Anselmo, A. C.; Mitragotri, S. The Evolution of Commercial Drug Delivery Technologies. Nat Biomed Eng 2021, 5 (9), 951–967. 10.1038/s41551-021-00698-w.

(4) Mann, J. L.; Maikawa, C. L.; Smith, A. A. A.; Grosskopf, A. K.; Baker, S. W.; Roth, G. A.; Meis, C. M.; Gale, E. C.; Liong, C. S.; Correa, S.; Chan, D.; Stapleton, L. M.; Yu, A. C.; Muir, B.; Howard, S.; Postma, A.; Appel, E. A. An Ultrafast Insulin Formulation Enabled by High-Throughput Screening of Engineered Polymeric Excipients. Sci. Transl. Med. 2020, 12 (550), eaba6676. 10.1126/scitranslmed.aba6676.

(5) Maikawa, C. L.; Chen, P. C.; Vuong, E. T.; Nguyen, L. T.; Mann, J. L.; d’Aquino, A. I.; Lal, R. A.; Maahs, D. M.; Buckingham, B. A.; Appel, E. A. Ultra-Fast Insulin–Pramlintide Co-Formulation for Improved Glucose Management in Diabetic Rats. Advanced Science 2021, 8 (21), 2101575. 10.1002/advs.202101575.

(6) Rao, V. A.; Kim, J. J.; Patel, D. S.; Rains, K.; Estoll, C. R. A Comprehensive Scientific Survey of Excipients Used in Currently Marketed, Therapeutic Biological Drug Products. Pharm Res 2020, 37 (10), 200. 10.1007/s11095-020-02919-4.

(7) Kerwin, B. A. Polysorbates 20 and 80 Used in the Formulation of Protein Biotherapeutics: Structure and Degradation Pathways. JPharmSci 2008, 97 (8), 2924–2935. 10.1002/jps.21190.

(8) Dubey, S.; Giovannini, R. Stability of Biologics and the Quest for Polysorbate Alternatives. Trends in Biotechnology 2021, 39 (6), 546–549. 10.1016/j.tibtech.2020.10.007.

(9) Fares, H. M.; Carnovale, M.; Tabouguia, M. O. N.; Jordan, S.; Katz, J. S. Novel Surfactant Compatibility with Downstream Protein Bioprocesses. JPharmSci 2023, 112 (7), 1811–1820. 10.1016/j.xphs.2023.04.011.

(10) Brosig, S.; Cucuzza, S.; Serno, T.; Bechtold-Peters, K.; Buecheler, J.; Zivec, M.; Germershaus, O.; Gallou, F. Not the Usual Suspects: Alternative Surfactants for Biopharmaceuticals. ACS Appl. Mater. Interfaces 2023, 15 (29), 34540–34553. 10.1021/acsami.3c05610.

(11) Wuchner, K.; Yi, L.; Chery, C.; Nikels, F.; Junge, F.; Crotts, G.; Rinaldi, G.; Starkey, J. A.; Bechtold-Peters, K.; Shuman, M.; Leiss, M.; Jahn, M.; Garidel, P.; de Ruiter, R.; Richer, S. M.; Cao, S.; Peuker, S.; Huille, S.; Wang, T.; Le Brun, V. Industry Perspective on the Use and Characterization of Polysorbates for Biopharmaceutical Products Part 1: Survey Report on Current State and Common Practices for Handling and Control of Polysorbates. Journal of Pharmaceutical Sciences 2022, 111 (5), 1280–1291. 10.1016/j.xphs.2022.02.009.

(12) Yang, K.; Hewarathna, A.; Geerlof-Vidavsky, I.; Rao, V. A.; Gryniewicz-Ruzicka, C.; Keire, D. Screening of Polysorbate-80 Composition by High Resolution Mass Spectrometry with Rapid H/D Exchange. Anal. Chem. 2019, 91 (22), 14649–14656. 10.1021/acs.analchem.9b03809.

(13) Emanuele, M.; Balasubramaniam, B. Differential Effects of Commercial-Grade and Purified Poloxamer 188 on Renal Function. Drugs R D 2014, 14 (2), 73–83. 10.1007/s40268-014-0041-0.

(14) Chen, W.; Ross, A.; Steinhuber, B.; Hoffmann, G.; Oltra, N. S.; Ravuri, S. K. K.; Bond, S.; Bell, C.; Kopf, R. The Development and Qualification of Liquid Adsorption Chromatography for Poloxamer 188 Characterization. Journal of Chromatography A 2021, 1652, 462353. 10.1016/j.chroma.2021.462353.

(15) Freire Haddad, H.; Burke, J. A.; Scott, E. A.; Ameer, G. A. Clinical Relevance of Pre-Existing and Treatment-Induced Anti-Poly(Ethylene Glycol) Antibodies. Regen Eng Transl Med 2022, 8 (1), 32–42. 10.1007/s40883-021-00198-y.

(16) Sellaturay, P.; Nasser, S.; Ewan, P. Polyethylene Glycol–Induced Systemic Allergic Reactions (Anaphylaxis). The Journal of Allergy and Clinical Immunology: In Practice 2021, 9 (2), 670–675. 10.1016/j.jaip.2020.09.029.

(17) Kanthe, A. D.; Carnovale, M. R.; Katz, J. S.; Jordan, S.; Krause, M. E.; Zheng, S.; Ilott, A.; Ying, W.; Bu, W.; Bera, M. K.; Lin, B.; Maldarelli, C.; Tu, R. S. Differential Surface Adsorption Phenomena for Conventional and Novel Surfactants Correlates with Changes in Interfacial mAb Stabilization. *Mol*. Pharmaceutics 2022, 19 (9), 3100–3113. 10.1021/acs.molpharmaceut.2c00152.

(18) Katz, J. S.; Nolin, A.; Yezer, B. A.; Jordan, S. Dynamic Properties of Novel Excipient Suggest Mechanism for Improved Performance in Liquid Stabilization of Protein Biologics. *Mol*. Pharmaceutics 2019, 16 (1), 282–291. 10.1021/acs.molpharmaceut.8b00984.

(19) Hanson, M. G.; Katz, J. S.; Ma, H.; Putterman, M.; Yezer, B. A.; Petermann, O.; Reineke, T. M. Effects of Hydrophobic Tail Length Variation on Surfactant-Mediated Protein Stabilization. *Mol*. Pharmaceutics 2020, 17 (11), 4302–4311. 10.1021/acs.molpharmaceut.0c00737.

(20) Liu, H.; Jin, Y.; Menon, R.; Laskowich, E.; Bareford, L.; Vilmorin, P. de; Kolwyck, D.; Yeung, B.; Yi, L. Characterization of Polysorbate 80 by Liquid Chromatography-Mass Spectrometry to Understand Its Susceptibility to Degradation and Its Oxidative Degradation Pathway. JPharmSci 2022, 111 (2), 323–334. 10.1016/j.xphs.2021.08.017.

(21) Wang, J.; Sun, H.; Yang, H.; Yang, R.; Zhu, X.; Guo, S.; Huang, Y.; Xu, Y.; Li, C.; Tu, J.; Sun, C. Dessecting the Toxicological Profile of Polysorbate 80 (PS80): Comparative Analysis of Constituent Variability and Biological Impact Using a Zebrafish Model. European Journal of Pharmaceutical Sciences 2024, 198, 106796. 10.1016/j.ejps.2024.106796.

(22) Li, X.; Wang, Z.; Zheng, B.; Wang, Y.; Zhang, J. Novel Strategy to Rapidly Profile and Identify Oxidized Species of Polysorbate 80 Using Ultra-High-Performance Liquid Chromatography Coupled with High-Resolution Mass Spectrometry. Anal. Chem. 2023, 95 (24), 9156–9163. 10.1021/acs.analchem.2c04956.

(23) Klich, J. H.; Kasse, C. M.; Mann, J. L.; Huang, Y.; d’Aquino, A. I.; Grosskopf, A. K.; Baillet, J.; Fuller, G. G.; Appel, E. A. Stable High-Concentration Monoclonal Antibody Formulations Enabled by an Amphiphilic Copolymer Excipient. Advanced Therapeutics 2023, 6 (1), 2200102. 10.1002/adtp.202200102.

(24) Maikawa, C. L.; Mann, J. L.; Kannan, A.; Meis, C. M.; Grosskopf, A. K.; Ou, B. S.; Autzen, A. A. A.; Fuller, G. G.; Maahs, D. M.; Appel, E. A. Engineering Insulin Cold Chain Resilience to Improve Global Access. Biomacromolecules 2021, 22 (8), 3386–3395. 10.1021/acs.biomac.1c00474.

(25) Chan, D.; Chien, J.-C.; Axpe, E.; Blankemeier, L.; Baker, S. W.; Swaminathan, S.; Piunova, V. A.; Zubarev, D. Yu.; Maikawa, C. L.; Grosskopf, A. K.; Mann, J. L.; Soh, H. T.; Appel, E. A. Combinatorial Polyacrylamide Hydrogels for Preventing Biofouling on Implantable Biosensors. Advanced Materials 2022, 34 (24), 2109764. 10.1002/adma.202109764.

(26) Tamasi, M. J.; Patel, R. A.; Borca, C. H.; Kosuri, S.; Mugnier, H.; Upadhya, R.; Murthy, N. S.; Webb, M. A.; Gormley, A. J. Machine Learning on a Robotic Platform for the Design of Polymer–Protein Hybrids. Advanced Materials 2022, 34 (30), 2201809. 10.1002/adma.202201809.

(27) Reis, M.; Gusev, F.; Taylor, N. G.; Chung, S. H.; Verber, M. D.; Lee, Y. Z.; Isayev, O.; Leibfarth, F. A. Machine-Learning-Guided Discovery of 19F MRI Agents Enabled by Automated Copolymer Synthesis. J. Am. Chem. Soc. 2021, 143 (42), 17677–17689. 10.1021/jacs.1c08181.

(28) Ruan, Z.; Li, S.; Grigoropoulos, A.; Amiri, H.; Hilburg, S. L.; Chen, H.; Jayapurna, I.; Jiang, T.; Gu, Z.; Alexander-Katz, A.; Bustamante, C.; Huang, H.; Xu, T. Population-Based Heteropolymer Design to Mimic Protein Mixtures. Nature 2023, 615 (7951), 251–258. 10.1038/s41586-022-05675-0.

(29) Panganiban, B.; Qiao, B.; Jiang, T.; DelRe, C.; Obadia, M. M.; Nguyen, T. D.; Smith, A. A. A.; Hall, A.; Sit, I.; Crosby, M. G.; Dennis, P. B.; Drockenmuller, E.; Olvera de la Cruz, M.; Xu, T. Random Heteropolymers Preserve Protein Function in Foreign Environments. Science 2018, 359 (6381), 1239–1243. 10.1126/science.aao0335.

(30) Jiang, T.; Hall, A.; Eres, M.; Hemmatian, Z.; Qiao, B.; Zhou, Y.; Ruan, Z.; Couse, A. D.; Heller, W. T.; Huang, H.; de la Cruz, M. O.; Rolandi, M.; Xu, T. Single-Chain Heteropolymers Transport Protons Selectively and Rapidly. Nature 2020, 577 (7789), 216–220. 10.1038/s41586-019-1881-0.

(31) Kumar, R.; Le, N.; Tan, Z.; Brown, M. E.; Jiang, S.; Reineke, T. M. Efficient Polymer-Mediated Delivery of Gene-Editing Ribonucleoprotein Payloads through Combinatorial Design, Parallelized Experimentation, and Machine Learning. ACS Nano 2020, 14 (12), 17626–17639. 10.1021/acsnano.0c08549.

(32) Dykeman-Bermingham, P. A.; Bogen, M. P.; Chittari, S. S.; Grizzard, S. F.; Knight, A. S. Tailoring Hierarchical Structure and Rare Earth Affinity of Compositionally Identical Polymers via Sequence Control. J. Am. Chem. Soc. 2024, jacs.4c00440. 10.1021/jacs.4c00440.

(33) Vincent, M. P.; Bobbala, S.; Karabin, N. B.; Frey, M.; Liu, Y.; Navidzadeh, J. O.; Stack, T.; Scott, E. A. Surface Chemistry-Mediated Modulation of Adsorbed Albumin Folding State Specifies Nanocarrier Clearance by Distinct Macrophage Subsets. Nat Commun 2021, 12 (1), 648. 10.1038/s41467-020-20886-7.

(34) Houang, E. M.; Haman, K. J.; Kim, M.; Zhang, W.; Lowe, D. A.; Sham, Y. Y.; Lodge, T. P.; Hackel, B. J.; Bates, F. S.; Metzger, J. M. Chemical End Group Modified Diblock Copolymers Elucidate Anchor and Chain Mechanism of Membrane Stabilization. *Mol*. Pharmaceutics 2017, 14 (7), 2333–2339. 10.1021/acs.molpharmaceut.7b00197.

(35) Gentekos, D. T.; Sifri, R. J.; Fors, B. P. Controlling Polymer Properties through the Shape of the Molecular-Weight Distribution. Nat Rev Mater 2019, 4 (12), 761–774. 10.1038/s41578-019-0138-8.

(36) Rosenbloom, S. I.; Hsu, J. H.; Fors, B. P. Controlling the Shape of the Molecular Weight Distribution for Tailored Tensile and Rheological Properties in Thermoplastics and Thermoplastic Elastomers. Journal of Polymer Science 2022, 60 (8), 1291–1299. 10.1002/pol.20210894.

(37) Gentekos, D. T.; Dupuis, L. N.; Fors, B. P. Beyond Dispersity: Deterministic Control of Polymer Molecular Weight Distribution. J. Am. Chem. Soc. 2016, 138 (6), 1848–1851. 10.1021/jacs.5b13565.

(38) Johnson, L. M.; Li, Z.; LaBelle, A. J.; Bates, F. S.; Lodge, T. P.; Hillmyer, M. A. Impact of Polymer Excipient Molar Mass and End Groups on Hydrophobic Drug Solubility Enhancement. Macromolecules 2017, 50 (3), 1102–1112. 10.1021/acs.macromol.6b02474.

(39) Ohnsorg, M. L.; Prendergast, P. C.; Robinson, L. L.; Bockman, M. R.; Bates, F. S.; Reineke, T. M. Bottlebrush Polymer Excipients Enhance Drug Solubility: Influence of End-Group Hydrophilicity and Thermoresponsiveness. ACS Macro Lett. 2021, 10 (3), 375–381. 10.1021/acsmacrolett.0c00890.

(40) Bianco, S.; Hasan, M.; Ahmad, A.; Richards, S.-J.; Dietrich, B.; Wallace, M.; Tang, Q.; Smith, A. J.; Gibson, M. I.; Adams, D. J. Mechanical Release of Homogenous Proteins from Supramolecular Gels. Nature 2024, 631 (8021), 544–548. 10.1038/s41586-024-07580-0.

(41) Liu, G. W.; Prossnitz, A. N.; Eng, D. G.; Cheng, Y.; Subrahmanyam, N.; Pippin, J. W.; Lamm, R. J.; Ngambenjawong, C.; Ghandehari, H.; Shankland, S. J.; Pun, S. H. Glomerular Disease Augments Kidney Accumulation of Synthetic Anionic Polymers. Biomaterials 2018, 178, 317–325. 10.1016/j.biomaterials.2018.06.001.

(42) Webber, M. J.; Appel, E. A.; Vinciguerra, B.; Cortinas, A. B.; Thapa, L. S.; Jhunjhunwala, S.; Isaacs, L.; Langer, R.; Anderson, D. G. Supramolecular PEGylation of Biopharmaceuticals. Proc Natl Acad Sci U S A 2016, 113 (50), 14189–14194. 10.1073/pnas.1616639113.

(43) Raffa, P.; Wever, D. A. Z.; Picchioni, F.; Broekhuis, A. A. Polymeric Surfactants: Synthesis, Properties, and Links to Applications. Chem. Rev. 2015, 115 (16), 8504–8563. 10.1021/cr500129h.

(44) Whitfield, R.; Truong, N. P.; Anastasaki, A. Precise Control of Both Dispersity and Molecular Weight Distribution Shape by Polymer Blending. Angewandte Chemie 2021, 133 (35), 19532–19537. 10.1002/ange.202106729.

(45) Whitfield, R.; Parkatzidis, K.; Truong, N. P.; Junkers, T.; Anastasaki, A. Tailoring Polymer Dispersity by RAFT Polymerization: A Versatile Approach. Chem 2020, 6 (6), 1340–1352. 10.1016/j.chempr.2020.04.020.

(46) Smith, A. A. A.; Hall, A.; Wu, V.; Xu, T. Practical Prediction of Heteropolymer Composition and Drift. ACS Macro Lett. 2019, 8 (1), 36–40. 10.1021/acsmacrolett.8b00813.

(47) Yu, H.; Liu, L.; Yin, R.; Jayapurna, I.; Wang, R.; Xu, T. Mapping Composition Evolution through Synthesis, Purification, and Depolymerization of Random Heteropolymers. J. Am. Chem. Soc. 2024, 146 (9), 6178–6188. 10.1021/jacs.3c13909.

(48) Smith, A. A. A.; Maikawa, C. L.; Hernandez, H. L.; Appel, E. A. Controlling Properties of Thermogels by Tuning Critical Solution Behaviour of Ternary Copolymers. Polym. Chem. 2021, 12 (13), 1918– 1923. 10.1039/D0PY01696A.

(49) Das, A.; Gangarde, Y. M.; Pariary, R.; Bhunia, A.; Saraogi, I. An Amphiphilic Small Molecule Drives Insulin Aggregation Inhibition and Amyloid Disintegration. International Journal of Biological Macromolecules 2022, 218, 981–991. 10.1016/j.ijbiomac.2022.07.155.

(50) Ivanova, M. I.; Sievers, S. A.; Sawaya, M. R.; Wall, J. S.; Eisenberg, D. Molecular Basis for Insulin Fibril Assembly. Proceedings of the National Academy of Sciences 2009, 106 (45), 18990–18995. 10.1073/pnas.0910080106.

(51) Chiang, H.-L.; Ngo, S. T.; Chen, C.-J.; Hu, C.-K.; Li, M. S. Oligomerization of Peptides LVEALYL and RGFFYT and Their Binding Affinity to Insulin. PLoS One 2013, 8 (6), e65358. 10.1371/journal.pone.0065358.

