## Supplementary Material for "Defining Structure-Function Relationships of Amphiphilic Excipients Enables Rational Design of Ultra-Stable Biopharmaceuticals"

#### Materials

All reagent grade materials and solvents were purchased from Sigma Aldrich, Tokyo Chemical Industry (TCI), or Fisher and used as received. Humalog (Insulin Lispro, Eli Lilly) was purchased and used as received. Slide-A-Lyzer dialysis cassettes (2 kDa MWCO) from thermofisher were used for polymer purification.

#### Instrumentation

##### Nuclear Magnetic Resonance (NMR) Spectroscopy

$^{1}$H NMR spectra were obtained using a Bruker Neo-500 MHz instrument. Solvents used in this study were obtained from Cambridge Isotope Laboratories and included: deuterium oxide and deuterated chloroform. Data was processed with MestReNova 10.0.

##### Size exclusion chromatography (SEC)

The molecular weight ($M_{n\ SEC}$) and dispersity were determined using an RI detector and polymethylmethacrylate standards. The running solvent was N,N-dimethyl formamide (DMF) with 1 g/L LiBr (flow rate: 1mL/min) heated to 50 °C and samples were prepared at 5 mg/mL. Separation was done through two Jordi Labs Resolve Mixed Bed Low Divinylbenzene (DVB) columns in series and data was collected by a Dionex Ultimate 3000 Variable Wavelength detector and RefractoMax521 RI detector. The RI traces were normalized and areas under the curves for the 310 nm absorbance signals were calculated with Prism 10.

##### Wilhelmy Plate

Surface tension at the air-water interface was measured with an Iridium/Platimum Wilhelmy plate connected to an electrobalance (KSV Nima, Finland). The set up was isolated in an environmental chamber to maintain humidity and temperature during the equilibration.

##### Plate Reader

Insulin aggregation was measured via non-specific absorbance at 540 nm in a BioTek Synergy H1 microplate reader.

##### Dynamic Light Scattering (DLS)

Self-assembly of amphiphilic polymers was assessed with a Wyatt DynaPro Plate Reader II. Samples were prepared in nanopure water at concentrations between 5 and 0.1 mg/mL and 5 measurements were taken and averaged per concentration.

### Methods

##### MoNi synthesis

Polymers were synthesized according to previously reported reversible addition fragmentation chain transfer (RAFT) polymerization procedures. As a representative example, 2-cyano-2-propyl dodecyl trithiocarbonate (CDPT, 82 mg, 0.24 mmol), 2,2′-azobis(2-methyl-propionitrile) (AIBN, 7.8 mg, 0.048 mmol), 4-acryloylmorpholine (Mo, 1.54g, 10.9 mmol), N-isopropylacrylamide (Ni, 460 mg, 4.07 mmol), and dimethyl formamide (DMF, 3.6 mL) were added to a 20 mL scintillation vial. To achieve various molecular weights only the CPDT and AIBN amounts were adjusted. The vial was capped with a PTFE septa and sparged for 20 minutes. The polymerization was conducted at 65 °C for 24 h to a conversion above 95% as determined by $H^1$ NMR. Polymers were purified by precipitating 3 times in 75:25 ether:hexane mixture with the exception of the 2 kDa MoNi which required a 50:50 mixture of ether and hexanes to precipitate. The materials were then dried under vacuum and stored at room temperature prior to end group removal.

##### HD polymer synthesis

To achieve high dispersity (HD) polymers with specific molecular weights, polymerization conditions from previously published work were adapted. Specifically, the CPDT was replaced with 2-Cyanopropan-2-yl N-methyl-N(pyridin-4-yl)carbamodithioate (CMCD). For example, to synthesize the 8 kDa HD MoNi, CMCD (29.8 mg, 0.0900 mmol), AIBN (3.8 mg, 0.023 mmol), Mo (560 mg, 3.96 mmol), Ni (167 mg, 1.48 mmol), and DMF (1.8 mL were added to a 20 mL scintillation vial. The reaction was purged for 1 h and polymerized at 65 °C for 16 h to a conversion of 95% via $H^1$ NMR. The polymer was purified by precipitation into 75:25 ether:hexanes. Removal of the CTA required 3 subsequent end-capping reactions to achieve >98% efficiency. In each reaction AIBN (656 mg, 4.0 mmol), lauryl peroxide (LPO, 159 mg, 0.4 mmol) and DMF (7.2 mL) were added to the polymer heated to 90 °C for 12 h. End group removal was measuring with UV absorbance at 310 nm during SEC and quantified by calculating the ratio of the areas under the curve (AUC) before and after end group removal. To tailor the polymer dispersity the HD polymers and low dispersity polymers were mixed in different ratios. For the 8 kDa MoNi mass ratios of low dispersity to HD polymers were 4, 1.5, 1, 0.667, 0.25, and 0 with dispersities of 1.20, 1.26, 1.29, 1.33, 1.40, and 1.62 as determined by SEC with PMMA standards. For the 4 kDa MoNi mass ratios were 3, 0.5, 0.33, and 0 with dispersities of 1.18, 1.24, 1.29, and 1.33 as determined by the same method.

##### End-group removal

Efficient end-group removal was required to yield polymers that could effectively act as excipients. We used the same excess radical end-group removal method (similar to previously published protocols) regardless of polymer molecular weight and radical initiator.[1,2] Briefly, polymers were precipitated and redissolved at 10 wt % in DMF. Then the radical initiator (AIBN, ACVA, VA-044, or V-70) at a 20-fold molar excess to the polymer chain ends and lauroyl peroxide (LPO) at a 2-fold molar excess was added to the solution. After sparging with nitrogen gas for 20 minutes, the solutions were sealed and heated to 90 °C for 24 h. The polymers were purified via precipitation into solutions of 75:25 ether:hexane 3 times (with the exception of the 2 kDa polymer which required a 50:50 mixture) and dried under vacuum. The dried samples were dissolved in nanopure water and dialyzed for 3 days and lyophilized to yield the final product. 2 kDa MWCO dialysis casettes were used for all polymers except the 2 kDa which was dialyzed with a 1 kDa MWCO dialysis membrane.

**MoDD synthesis**

As a representative example for MoDD10, 4-((((2-carboxyethyl)thio)carbonothioyl)thio)-4-cyanopentanoic acid (CCC, 16.4 mg, 0.54 mmol), 2,2′-azobis(2-methyl-propionitrile) (AIBN, 1.76 mg, 0.011 mmol), 4-acryloylmorpholine (Mo, 0.500 g, 3.55 mmol), N-dodecylacrylamide (DD, 62 mg, 0.26 mmol), and dimethyl formamide (DMF, 0.8 mL) were added to a 7 mL scintillation vial. The vial was capped with a rubber septa and sparged for 20 minutes. The polymerization was conducted at 65 °C for 24 h to a conversion of 98% as determined by $H^1$ NMR. Polymers were purified by precipitating 3 times in 75:25 ether:hexane mixture. The materials were then dried under vacuum, dialyzed with a 2 kDa MWCO membrane against DI water for 3 days and lyophilized.

### Experimental

#### Insulin stability

Accelerated aging studies to measure insulin stability were replicated from Webber et al. and Mann et al. with slight modifications.[2–4] Commercial Humalog was converted to monomeric insulin via the addition of EDTA (60 uL of 20 mM EDTA per 1 mL of Humalog). Polymers were dissolved at 100 mg/mL in nanopure water and to a 1 mL of monomeric insulin 2 uL was added to achieve a final concentration of 0.02 wt %. 200 uL of the insulin samples with and without MoNi were added to each well of a clear 96-well plate (in quadruplicate) and sealed with optically transparent plate covers. The plates were heated to 37 °C and shaken orbitally for up to 72 h, with absorbance measurements at 540 nm occurring every 10 minutes. Formation of amyloid fibrils causes occlusion of the clear wells which leads to high absorbance values. T10 Aggregation times were determined by normalizing absorbance values and assessing the time at which the threshold of 10% aggregation was exceeded. All stability measurements were done with 4 technical replicates and data plotted as mean and SEM.

#### Surface tension measurements

Equilibrium surface tension values and equilibration rate constants were determined with a Wilhelmy plate as previously reported.[5] To summarize, a simulated solution of Humalog buffer was made by adding, to 1 L of nanopure water, 16 g glycerin and 1.88 g dibasic sodium phosphate and was used for all surface tension measurements. To 10 mL of Humalog buffer, polymer was added to achieve a final concentration of 0.02 wt %. Time-resolved surface tension measurements were performed by partial immersion of a Wilhelmy plate and recorded with an electrobalance. Data was recorded over ~1.3 h (5000 s) to allow the system to reach equilibrium and avoid evaporation effects that occurred during experiments longer than 1.5 h. Experiments were repeated in duplicate.

#### Compositional Drift

The composition of MoNi polymer chains was based on the Mayo-Lewis model and simulated via Monte Carlo method according to previously published protocols.[6,6,7] New analysis and graphical representations of the distribution of monomer compositions was performed using novel open-source software (https://github.com/stetef/Compositional-Drift-with-Compositional-Dispersity-Index). To generate the specific data for this study, pool sizes of 200,000, chain transfer percentages (to simulate dispersity measured by SEC), degrees of polymerization (15 -150), and reactivity ratios (Morph 1.21 and Nipam 0.84, as determined in DMF) were used as simulation inputs.

#### Supplementary Figures and Tables

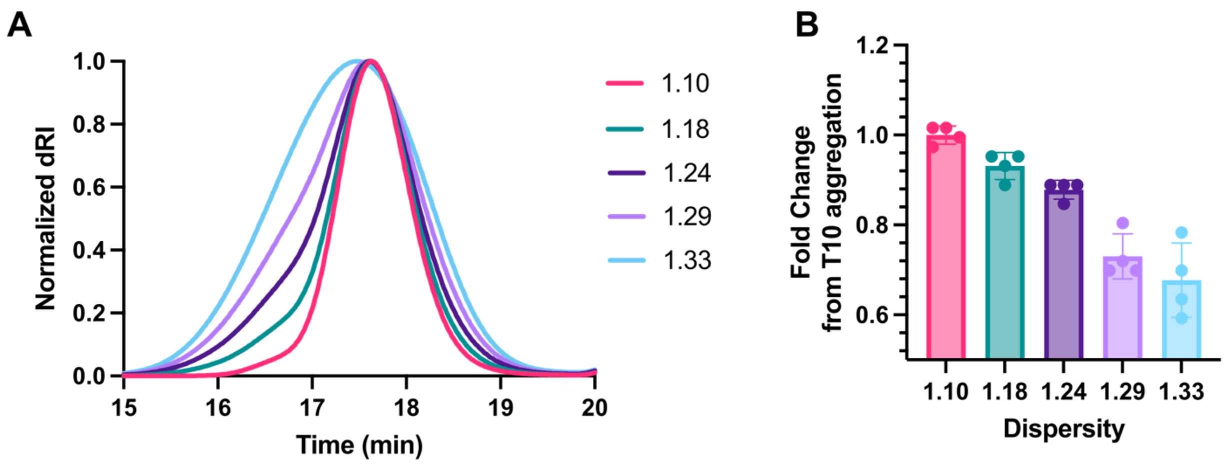

**Figure S1.** A) SEC chromatograms for mixtures of low and high dispersity 4 kDa $MoNi_{23\%}$ with identical compositions. B) Comparative T10 aggregation times relative to low dispersity 4 kDa $MoNi_{23\%}$ as a function dispersity.

**Table S1.** Evaluation of dispersity of $MoNi_{23\%}$ 4 and 4-HD mixtures with SEC and PMMA standards. End-group removal was determined by ratio of the area under the curve (AUC) before and after the reaction at 310 nm.

| Label | $M_n$ | $M_w$ | Dispersity | End group removal |
|---|---|---|---|---|
| 4 | 4,676 | 5,152 | 1.10 | 99.2% |
| 1.18 | 4,843 | 5,730 | 1.18 | --- |
| 1.24 | 5,011 | 6,232 | 1.24 | --- |
| 1.29 | 5,139 | 6,619 | 1.29 | --- |
| 4-HD | 5,279 | 6,996 | 1.33 | 93.06% |

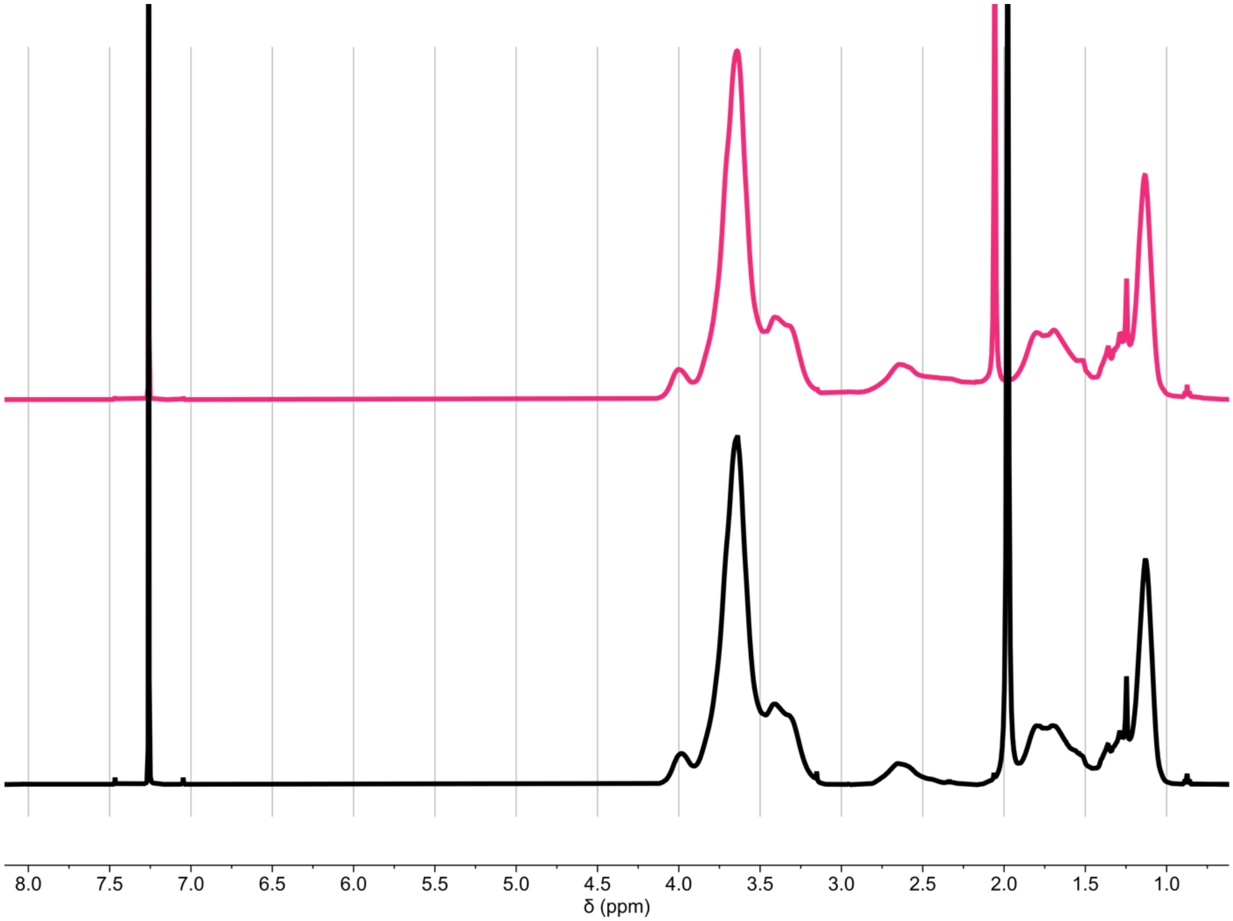

**Figure S2.** $^{1}$H NMR characterization of purified $MoNi_{23\%}$ polymers 4 kDa (top, —) and 4 kDa HD (bottom, —) in $CDCl_3$.

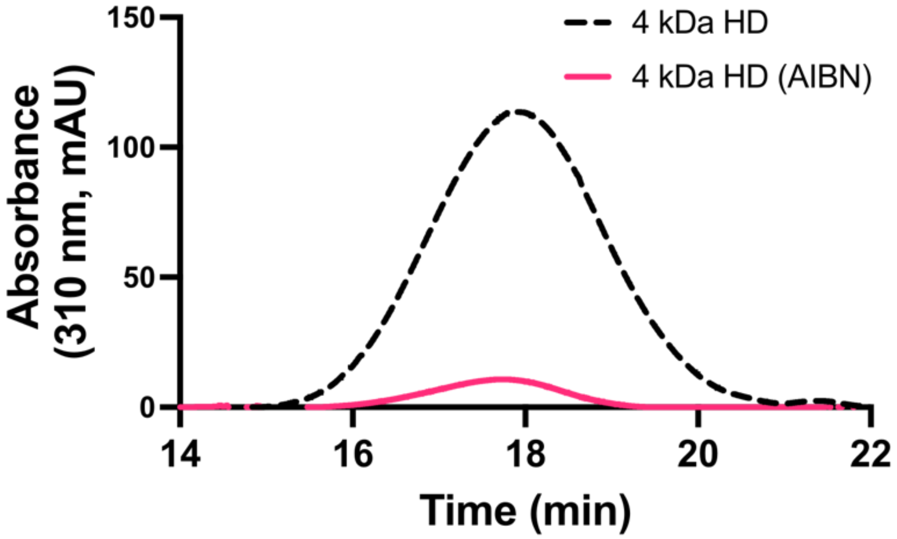

**Figure S3.** Quantification of CTA end-group removal via SEC and absorbance measurements. Measuring AUC of the peaks gives an end-group removal efficiency of 93.06%

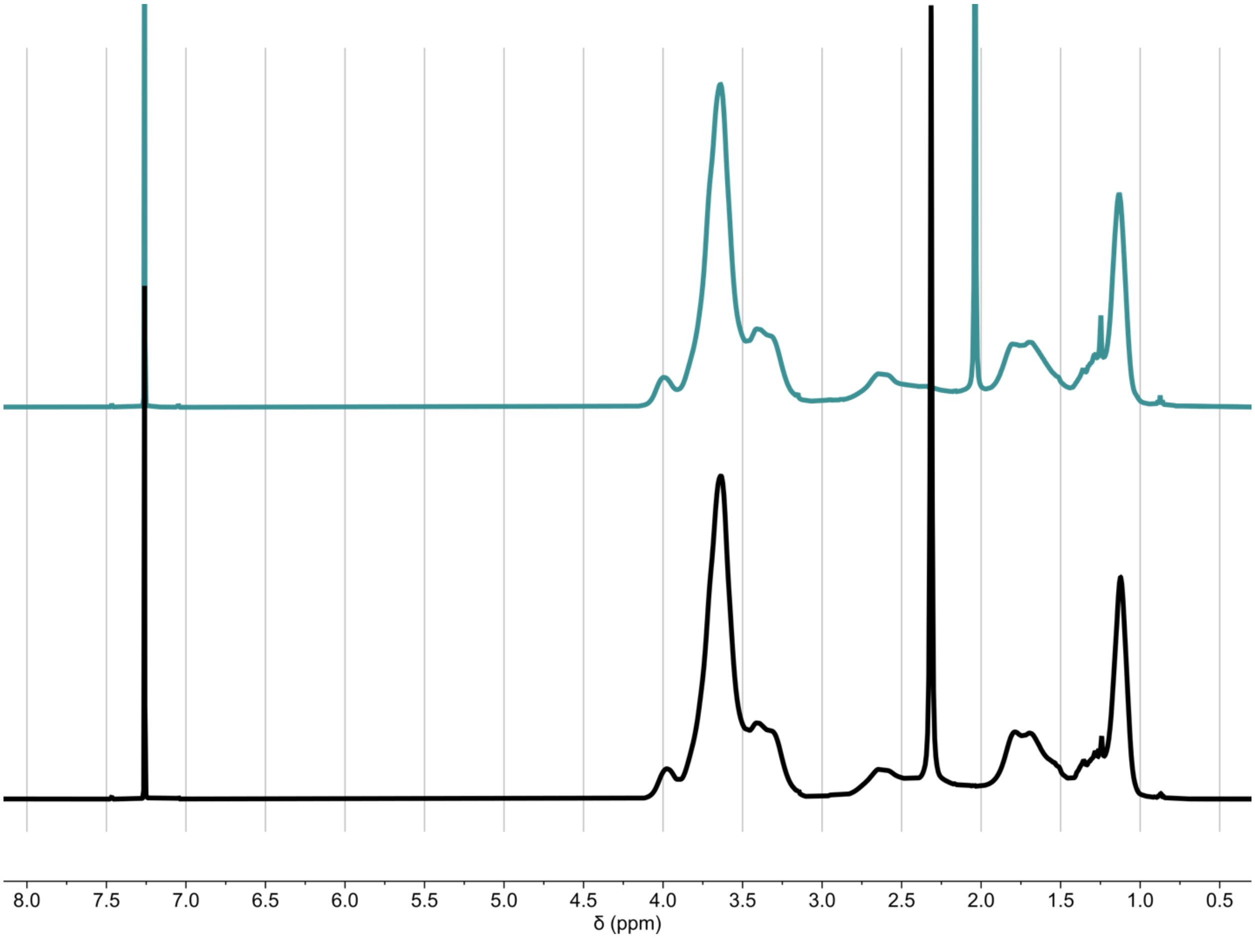

**Figure S4.** $^{1}$H NMR characterization of purified $MoNi_{23\%}$ polymers 8 kDa (top, —) and 8 kDa HD (bottom, —) in $CDCl_3$.

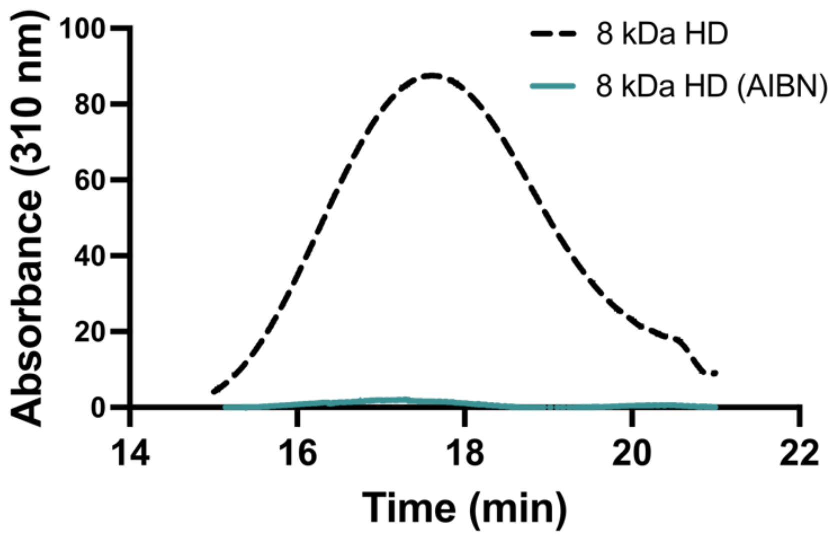

**Figure S5.** Quantification of CTA end-group removal via SEC and absorbance measurements. Measuring AUC of the peaks gives an end-group removal efficiency of 98.47%

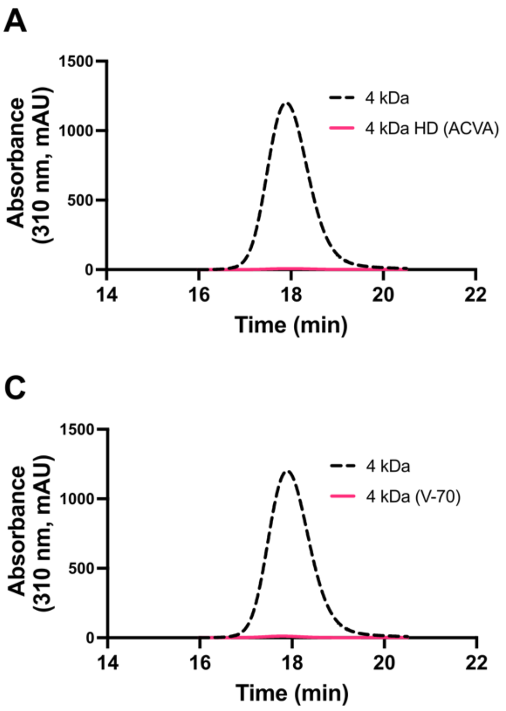

**Figure S6.** Quantification of CTA end-group removal via SEC and absorbance (310 nm) measurements with various thermal radical initiators.

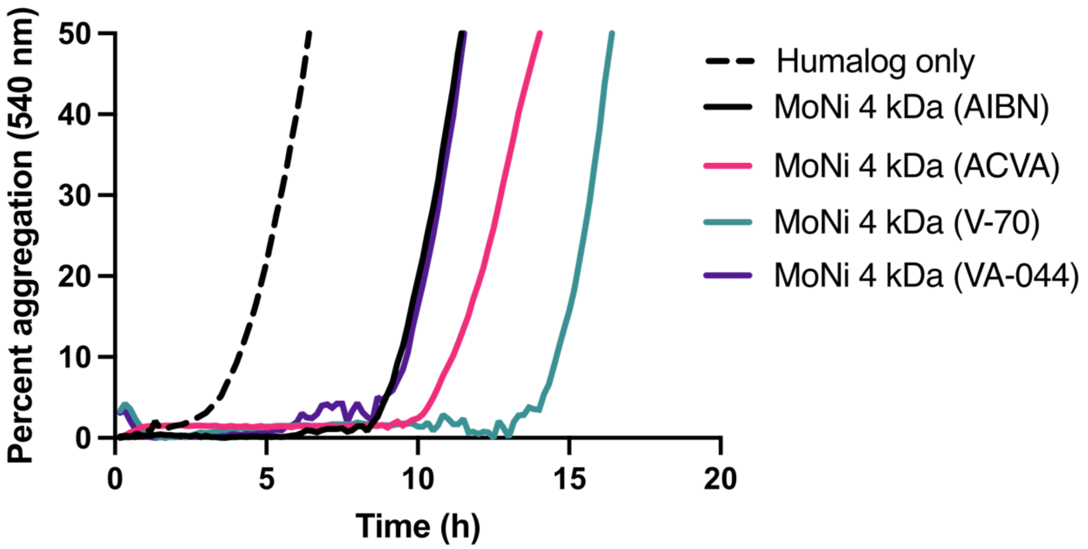

**Figure S7.** Non-specific absorbance assay to assess stability under stressed aging conditions (37 ˚C and shaking) of monomeric insulin with EDTA (1.2 mM) + polymer excipients (0.02 wt %) and commercial insulin hexamer (Humalog). Data presented are a mean of 4 technical replicates.

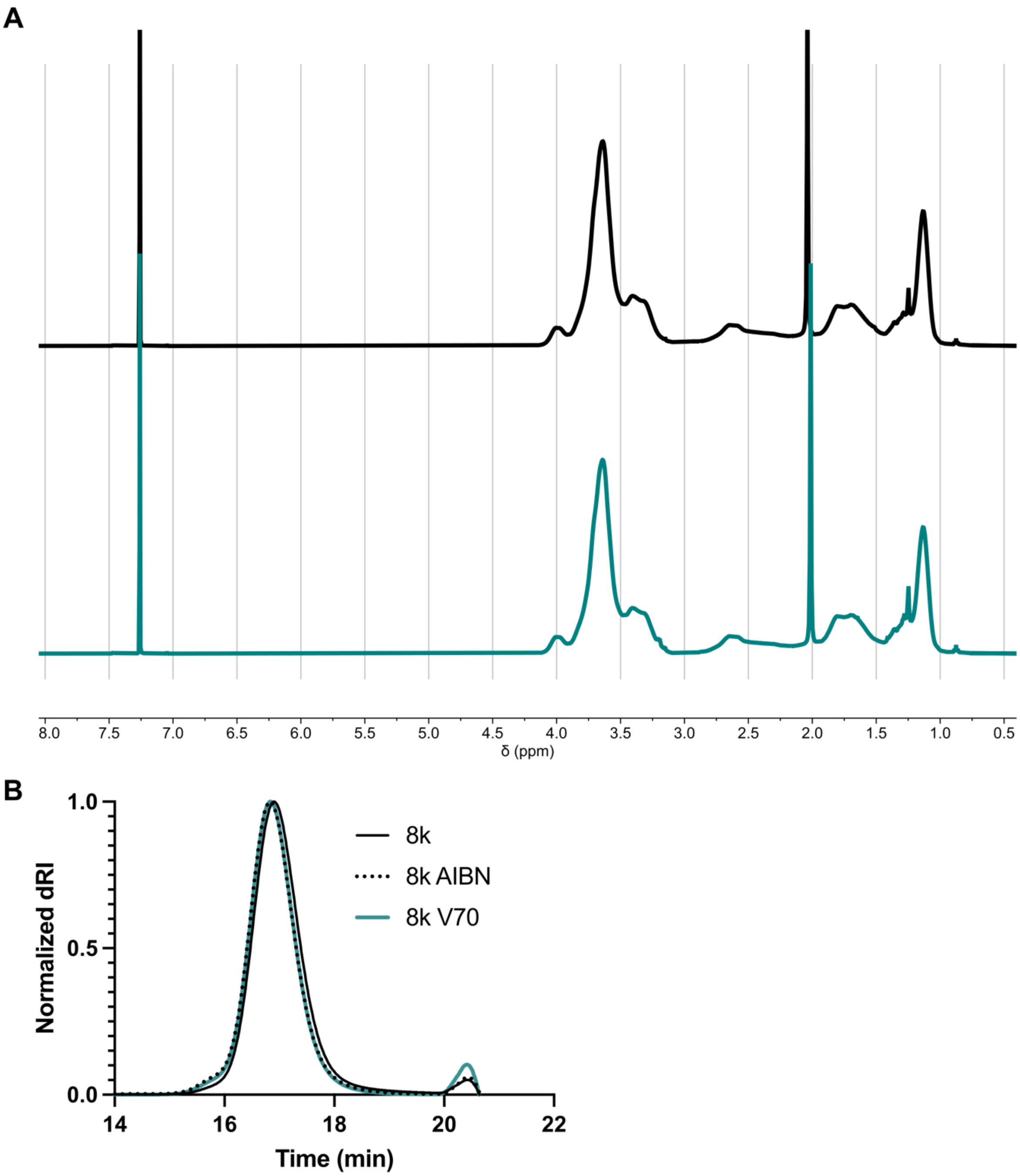

**Figure S8.** A) . [1]H NMR characterization of purified $MoNi_{23\%}$ polymers 8 kDa AIBN (top, —) and 8 kDa V70 (bottom, —) in $CDCl_3$. B) SEC chromatograms for 8 kDa with CTA (**—**), 8 kDa with AIBN end-group (**---**) and 8 kDa V70 end-group (—) $MoNi_{23\%}$ with identical compositions.

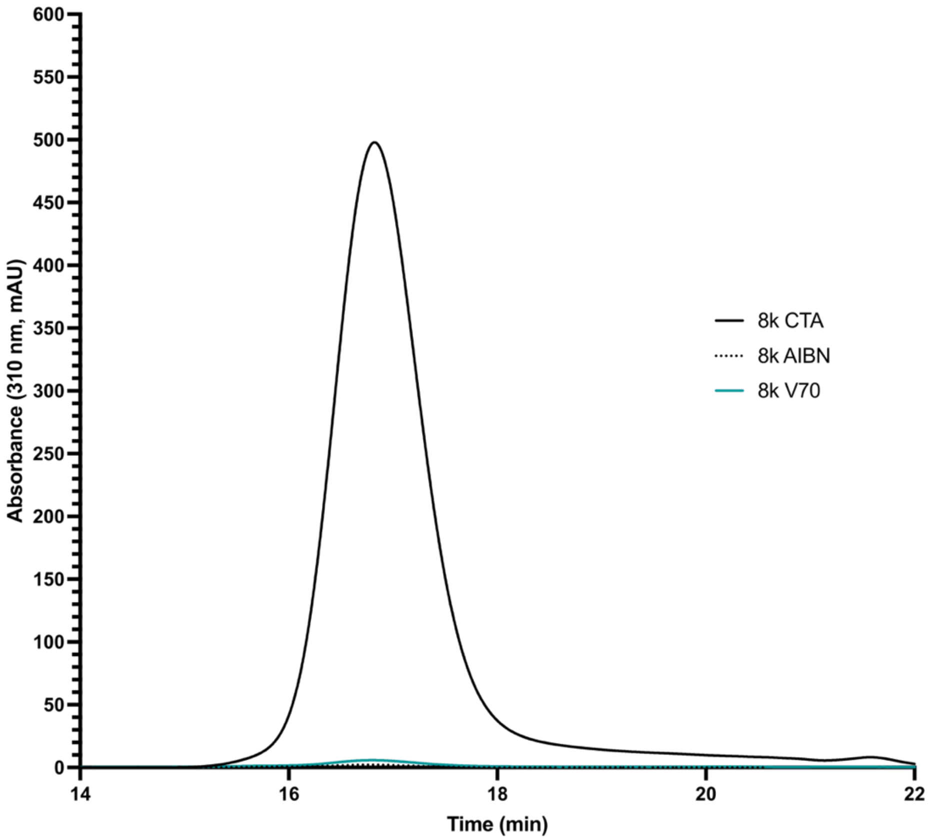

**Figure S9.** Quantification of CTA end-group removal via SEC and absorbance (310 nm) measurements for 8 kDa $MoNi_{23\%}$ with thermal radical initiators (AIBN and V70).

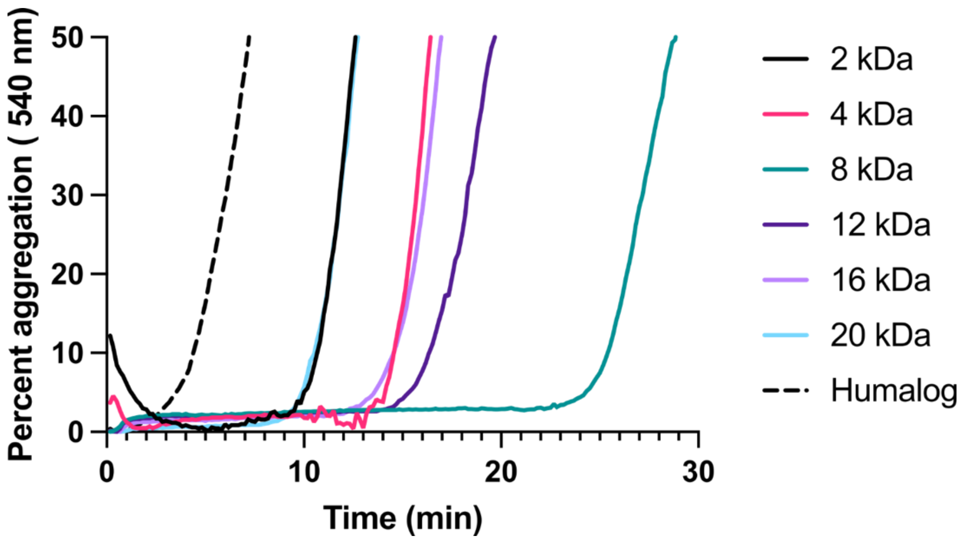

**Figure S10.** Non-specific absorbance assay to assess stability under stressed aging conditions (37 ˚C and shaking) of monomeric insulin with EDTA (1.2 mM) + polymer excipients with V-70 end-groups (0.02 wt %) and commercial insulin hexamer (Humalog). Data presented are a mean of 4 technical replicates.

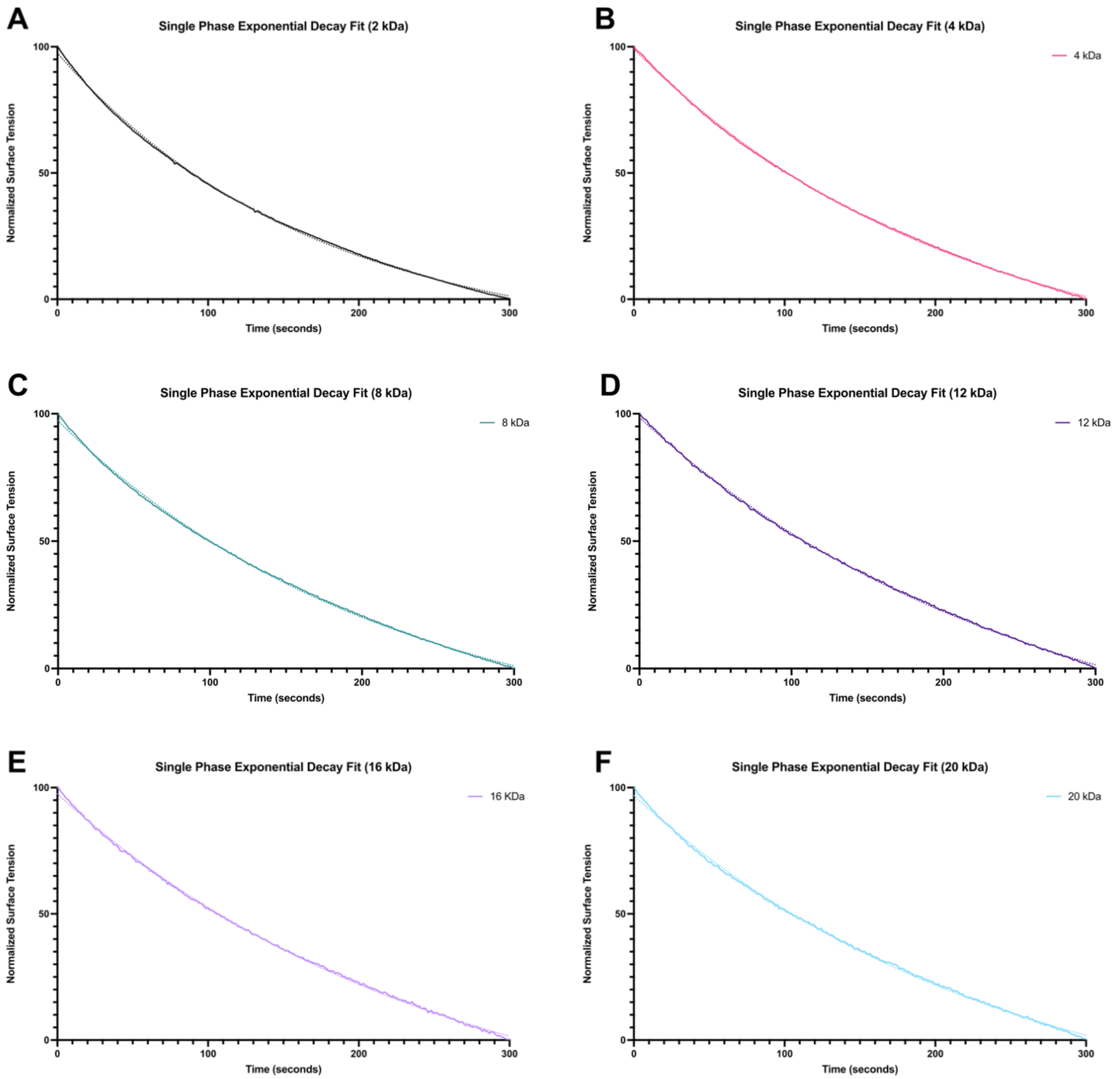

**Figure S11.** A-F) Curve fitting of the first 300 s (5 min) of time-resolved surface tensions measurements for 0.02 wt % solutions of $MoNi_{23\%}$ with various molecular weights in simulated Humalog buffer. Experimental data is presented as the solid line and fits as the dotted lines. All fits were performed with Prism 10.

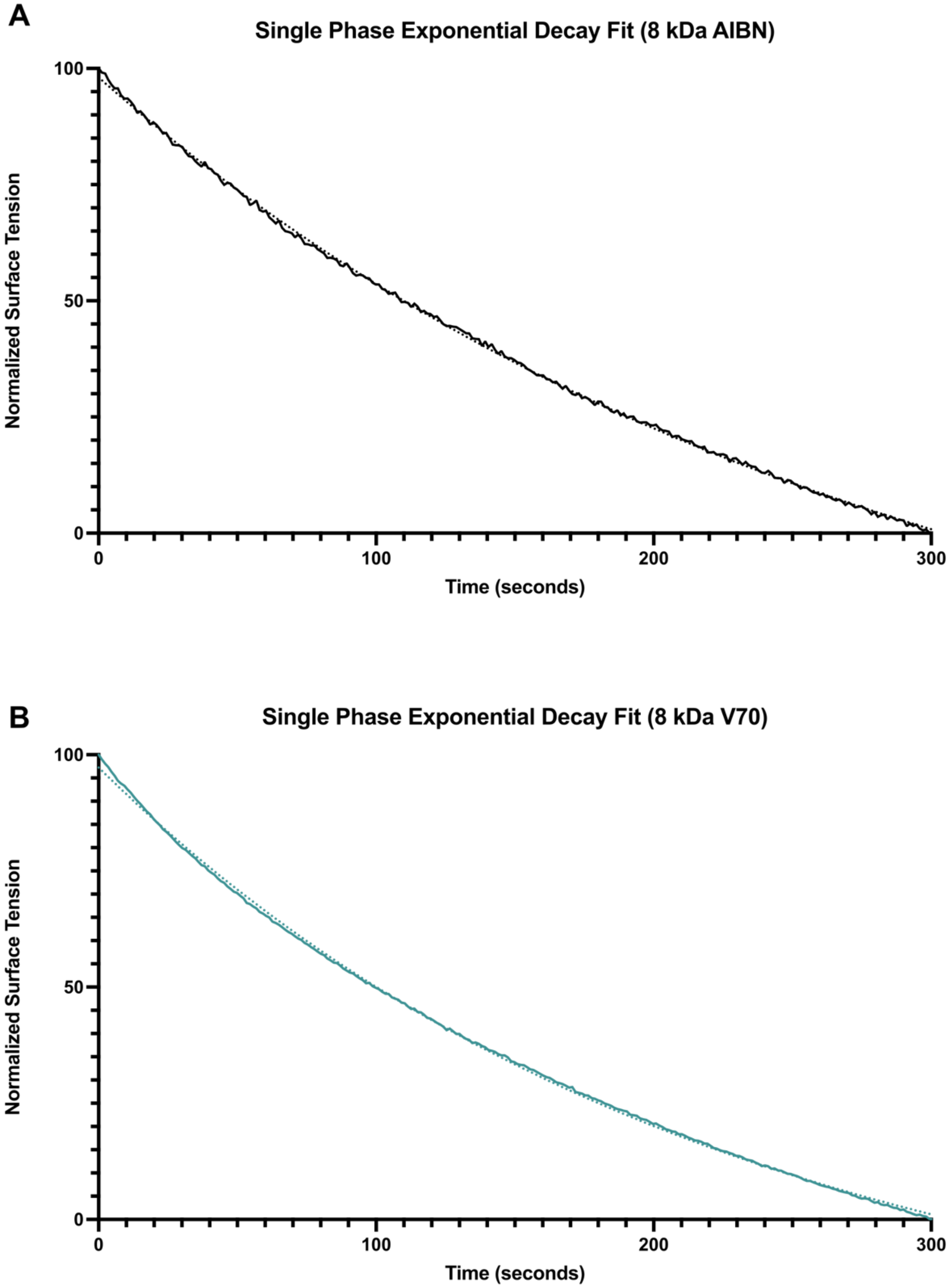

**Figure S12.** A) Curve fitting of the first 300 s (5 min) of time-resolved surface tensions measurements for 0.02 wt % solutions of $MoNi_{23\%}$ with 8 kDa and AIBN end-group in simulated Humalog buffer. B) Curve fitting of the first 300 s (5 min) of time-resolved surface tensions measurements for 0.02 wt % solutions of $MoNi_{23\%}$ with 8 kDa and V-70 end-group in simulated Humalog buffer. Experimental data is presented as the solid line and fits as the dotted lines. All fits were performed with Prism 10.

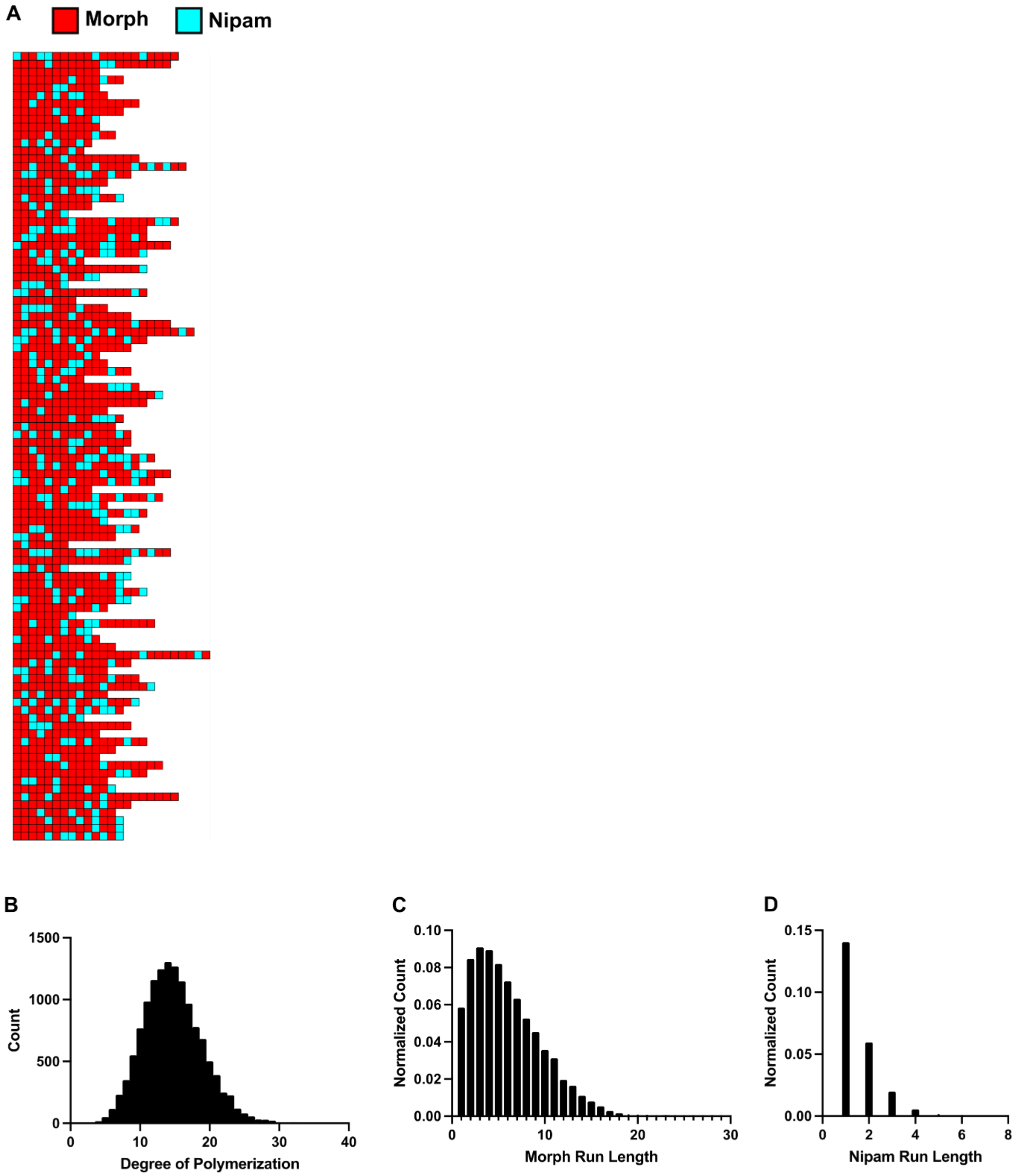

**Figure S13.** Simulation data from Compositional Drift for 2 kDa $MoNi_{23\%}$. A) 100 example chains out of 200,000 demonstrating monomer distributions. B) Size distribution of generated polymer chains showing dispersity. C) Distribution of number of Morph monomers in a continuous block. D) Distribution of number of Nipam monomers in a continuous block.

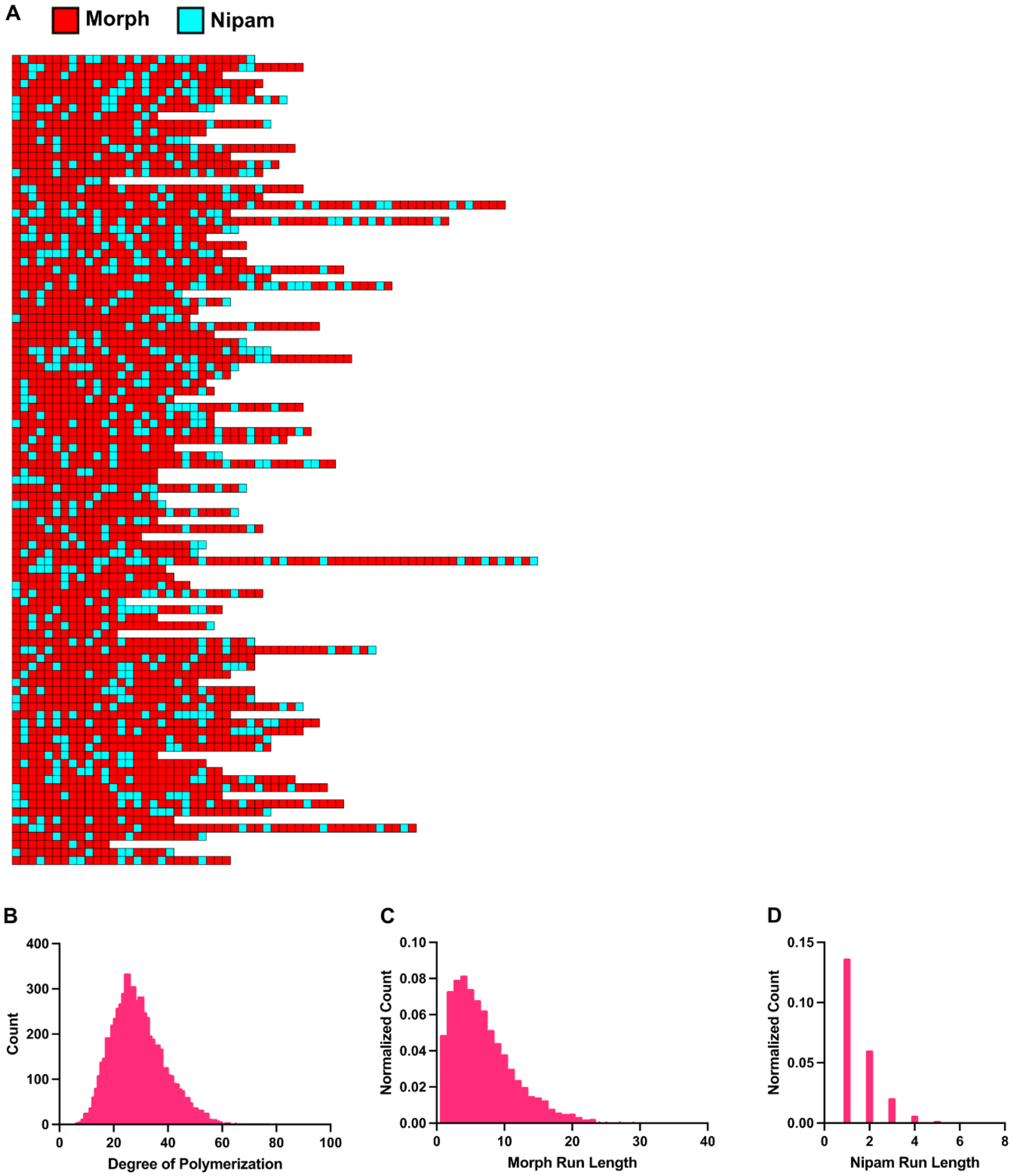

**Figure S14.** Simulation data from Compositional Drift for 4 kDa $MoNi_{23\%}$. A) 100 example chains out of 200,000 demonstrating monomer distributions. B) Size distribution of generated polymer chains showing dispersity. C) Distribution of number of Morph monomers in a continuous block. D) Distribution of number of Nipam monomers in a continuous block.

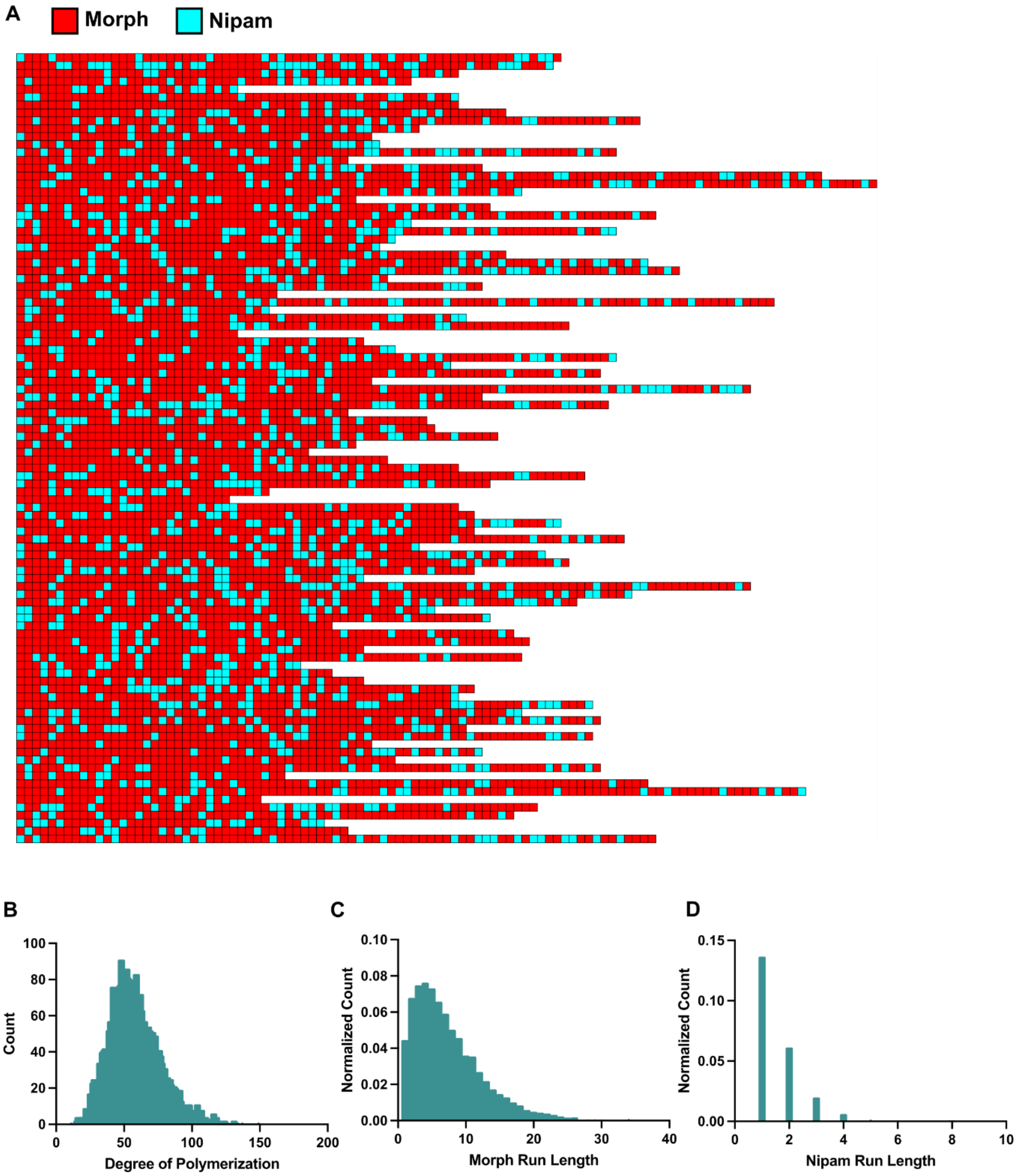

**Figure S15.** Simulation data from Compositional Drift for 8 kDa $MoNi_{23\%}$. A) 100 example chains out of 200,000 demonstrating monomer distributions. B) Size distribution of generated polymer chains showing dispersity. C) Distribution of number of Morph monomers in a continuous block. D) Distribution of number of Nipam monomers in a continuous block.

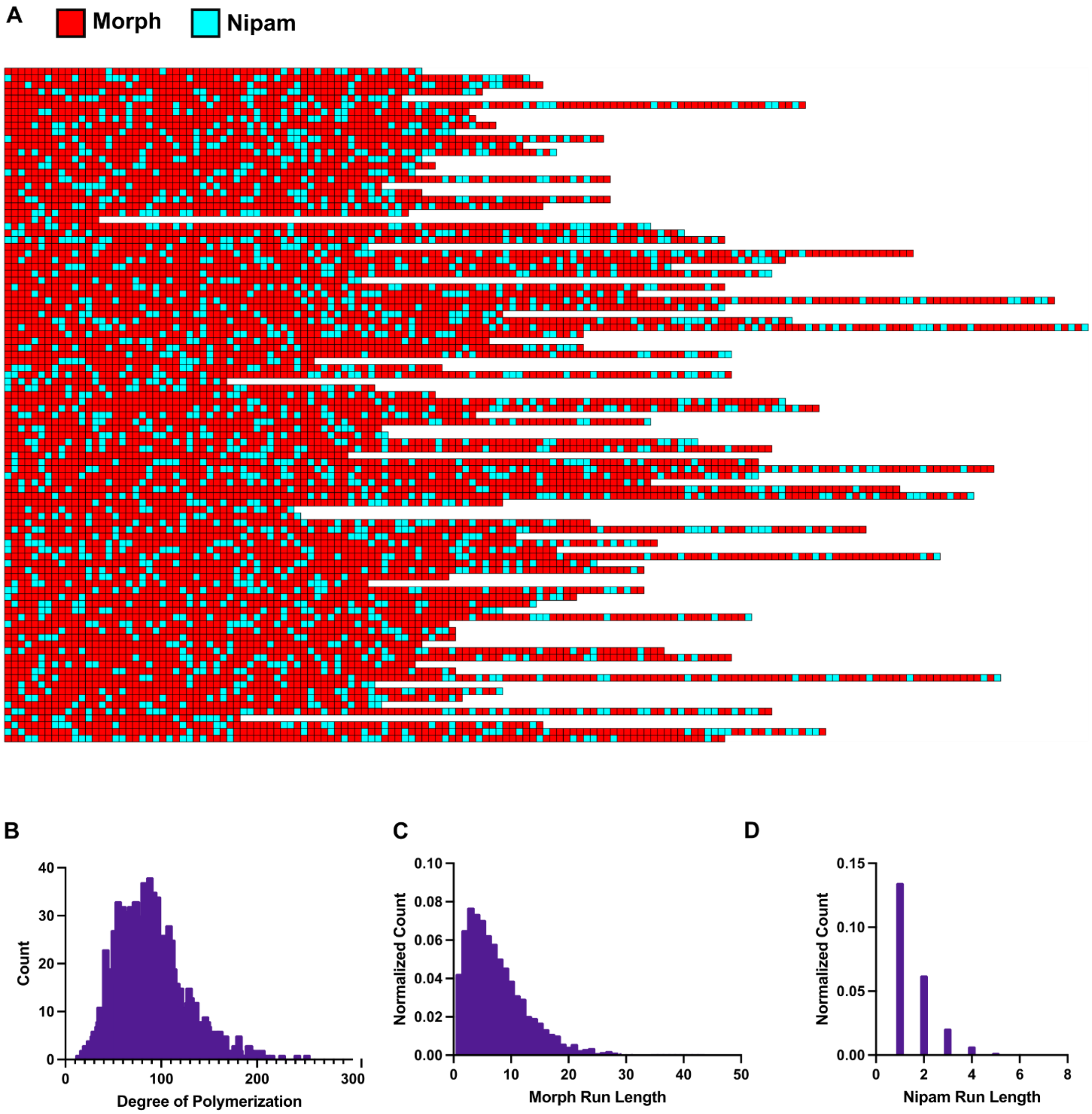

**Figure S16.** Simulation data from Compositional Drift for 12 kDa $MoNi_{23\%}$. A) 100 example chains out of 200,000 demonstrating monomer distributions. B) Size distribution of generated polymer chains showing dispersity. C) Distribution of number of Morph monomers in a continuous block. D) Distribution of number of Nipam monomers in a continuous block.

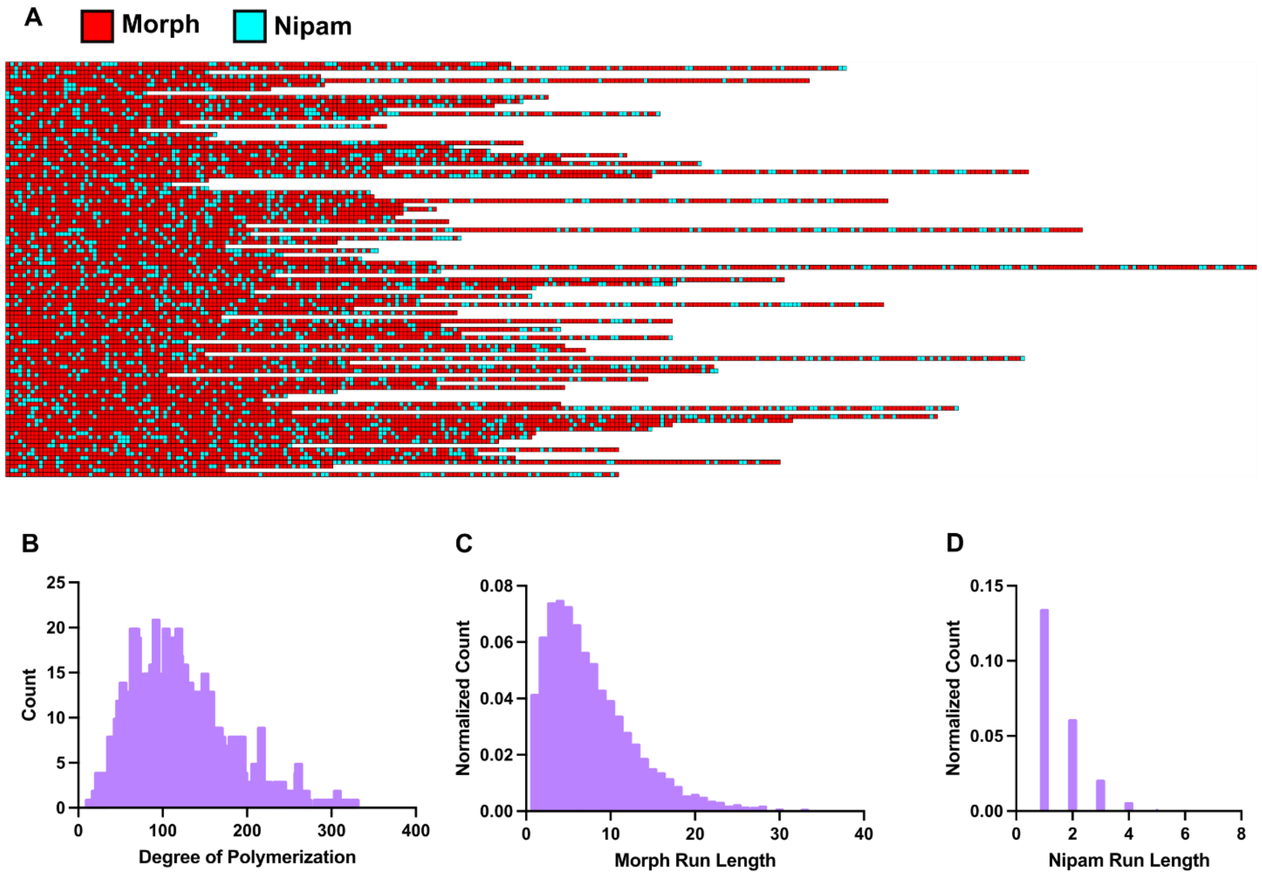

**Figure S17.** Simulation data from Compositional Drift for 16 kDa $MoNi_{23\%}$. A) 100 example chains out of 200,000 demonstrating monomer distributions. B) Size distribution of generated polymer chains showing dispersity. C) Distribution of number of Morph monomers in a continuous block. D) Distribution of number of Nipam monomers in a continuous block.

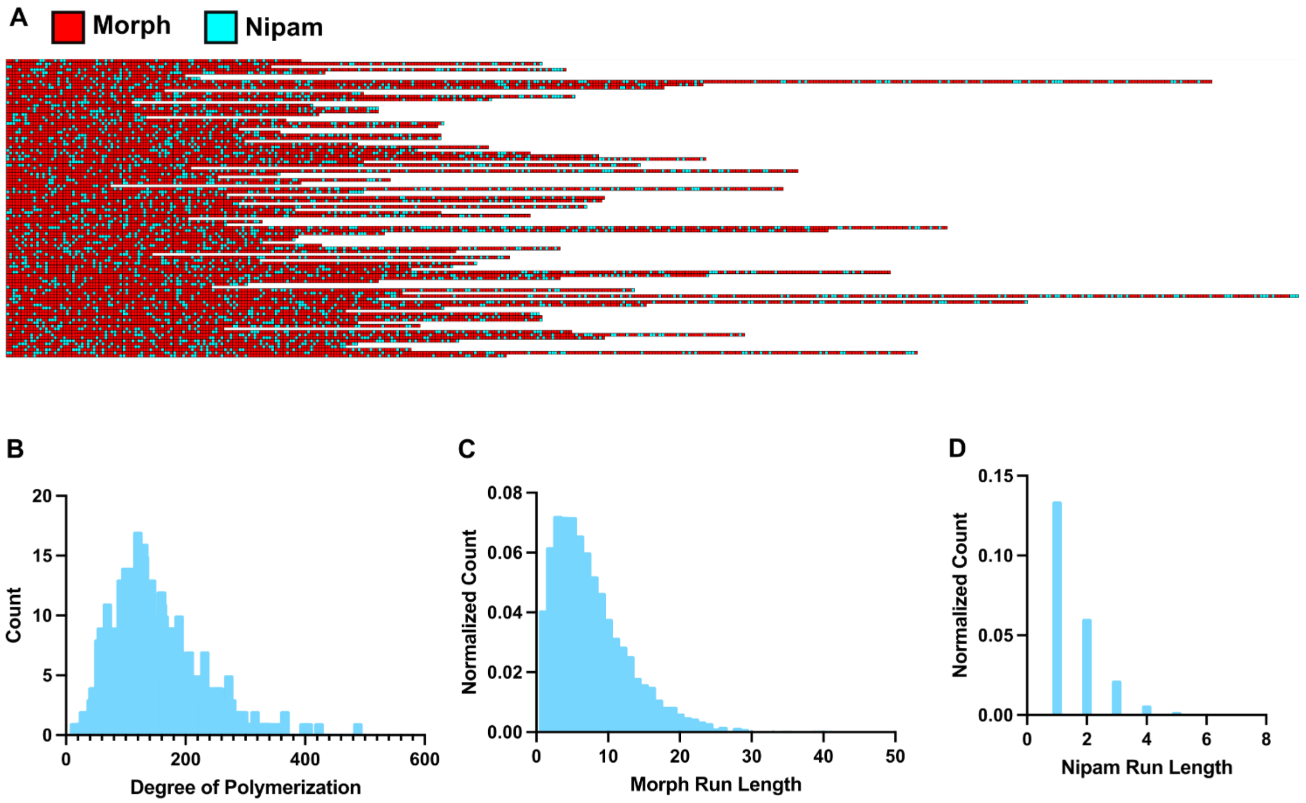

**Figure S18.** Simulation data from Compositional Drift for 20 kDa $MoNi_{23\%}$. A) 100 example chains out of 200,000 demonstrating monomer distributions. B) Size distribution of generated polymer chains showing dispersity. C) Distribution of number of Morph monomers in a continuous block. D) Distribution of number of Nipam monomers in a continuous block.

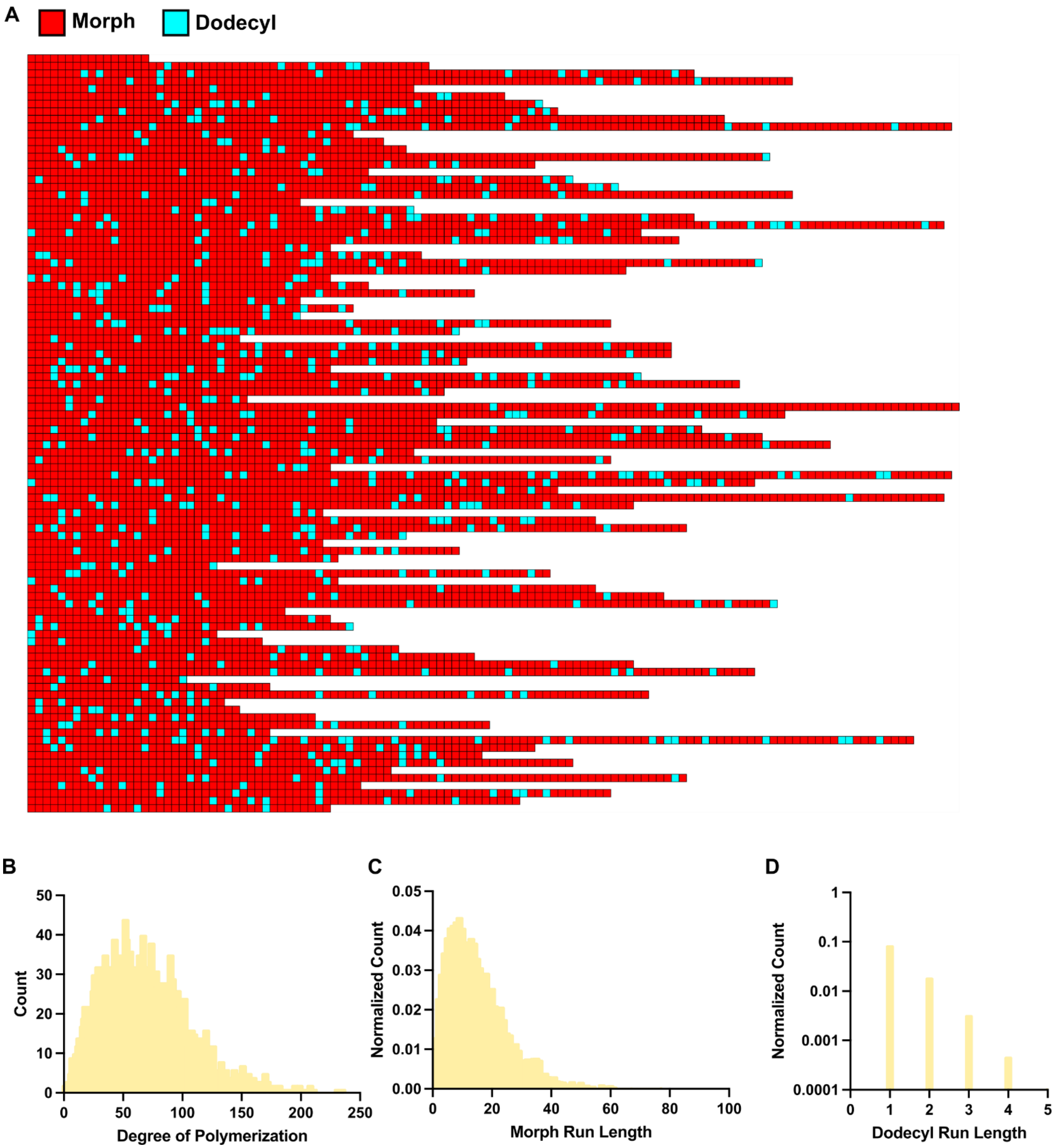

**Figure S19.** Simulation data from Compositional Drift for 10 kDa MoDD. A) 100 example chains out of 200,000 demonstrating monomer distributions. B) Size distribution of generated polymer chains showing dispersity. C) Distribution of number of Morph monomers in a continuous block. D) Distribution of number of Dodecyl monomers in a continuous block.

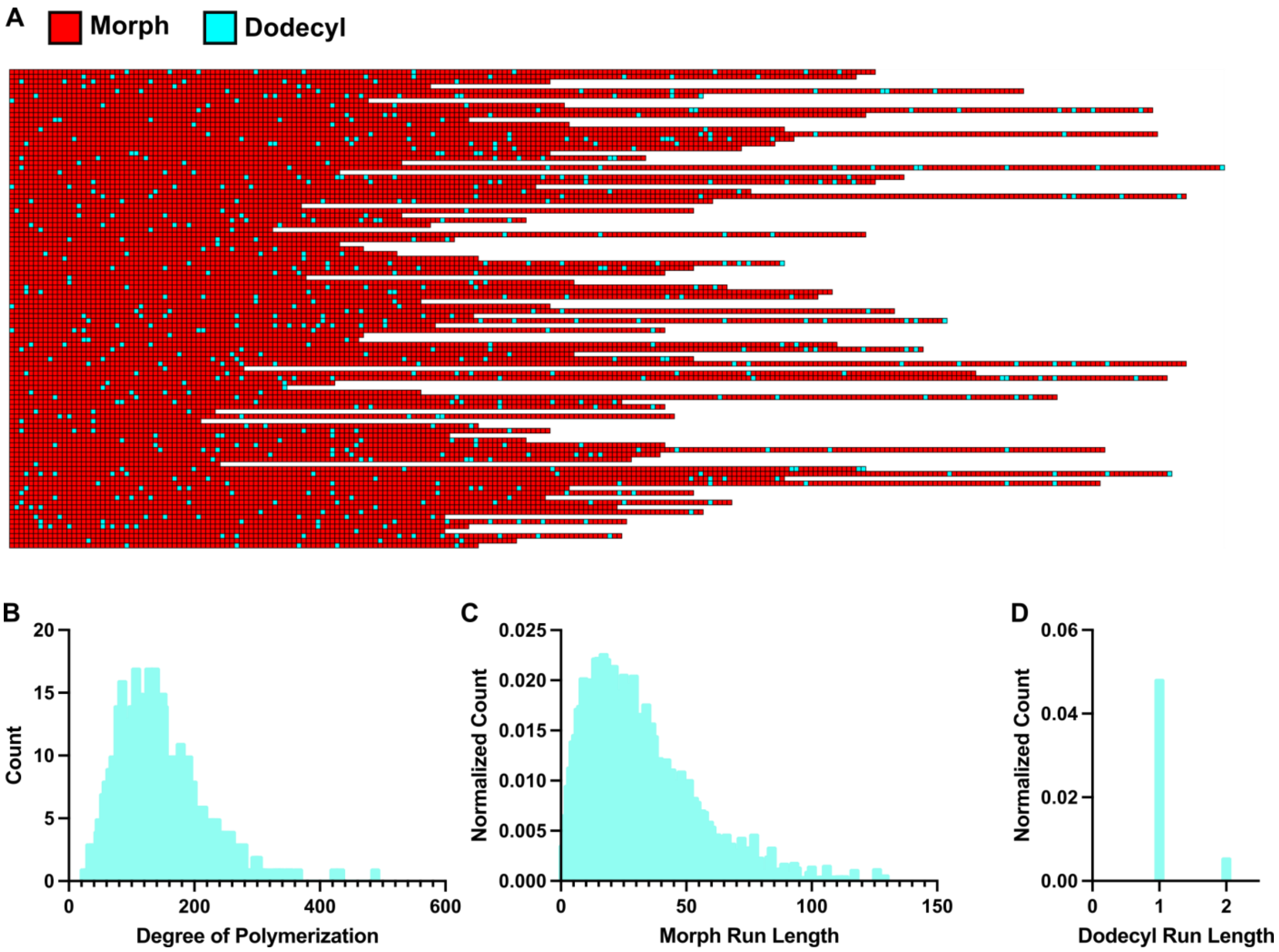

**Figure S20.** Simulation data from Compositional Drift for 20 kDa MoDD. A) 100 example chains out of 200,000 demonstrating monomer distributions. B) Size distribution of generated polymer chains showing dispersity. C) Distribution of number of Morph monomers in a continuous block. D) Distribution of number of Dodecyl monomers in a continuous block.

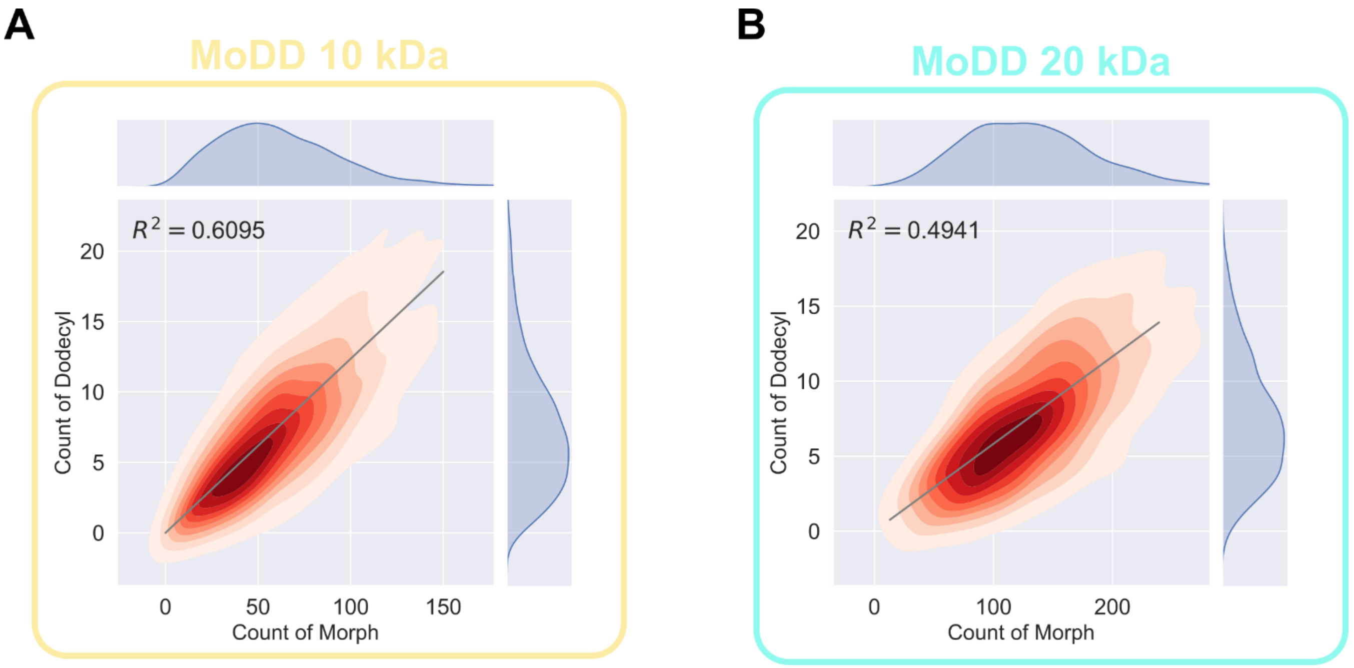

**Figure S21.** 2D-plots of incorporation of Morph and Dodecyl simulated for individual polymer chains to visualize monomer compositions of MoDD polymers with various molecular weights and monomer feed ratios. A) To quantify deviation from the ideal monomer composition for MoDD10 based on feed ratio (Morph/Dodecyl of 89/11), $R^2$ was calculated for the distributions against a line with a slope of 11/89. B) To quantify deviation from the ideal monomer composition for MoDD20 based on feed ratio (Morph/Dodecyl of 94.5/5.5), $R^2$ was calculated for the distributions against a line with a slope of 5.5/94.5.

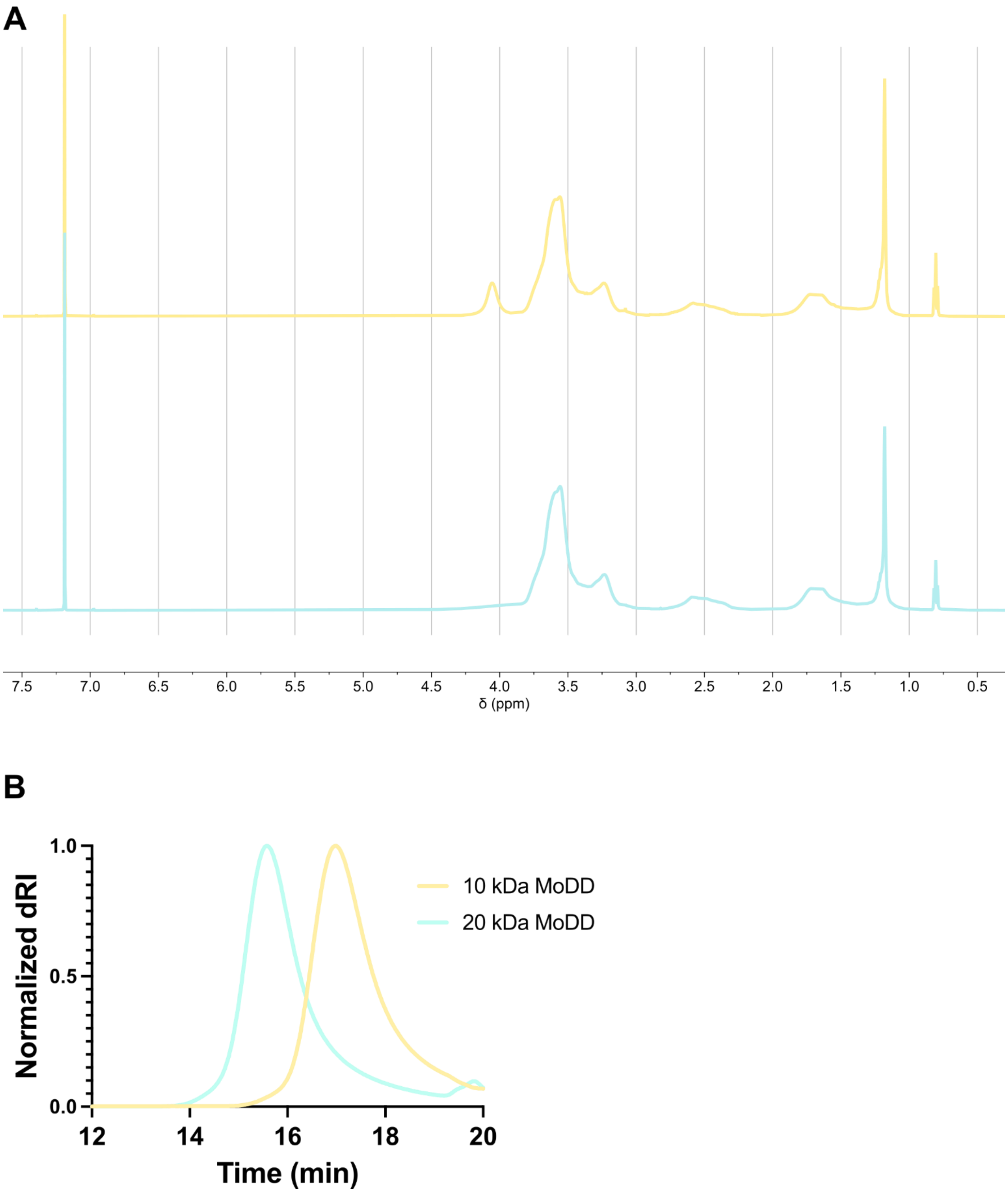

**Figure S22.** A) [1]H NMR characterization of purified MoDD polymers of various molecular weights (top, —, 10 kDa and bottom, —, 20 kDa) in $CDCl_3$. C) Size exclusion chromatography (SEC) determination of MoDD molecular weight and dispersity relative to PMMA standards.

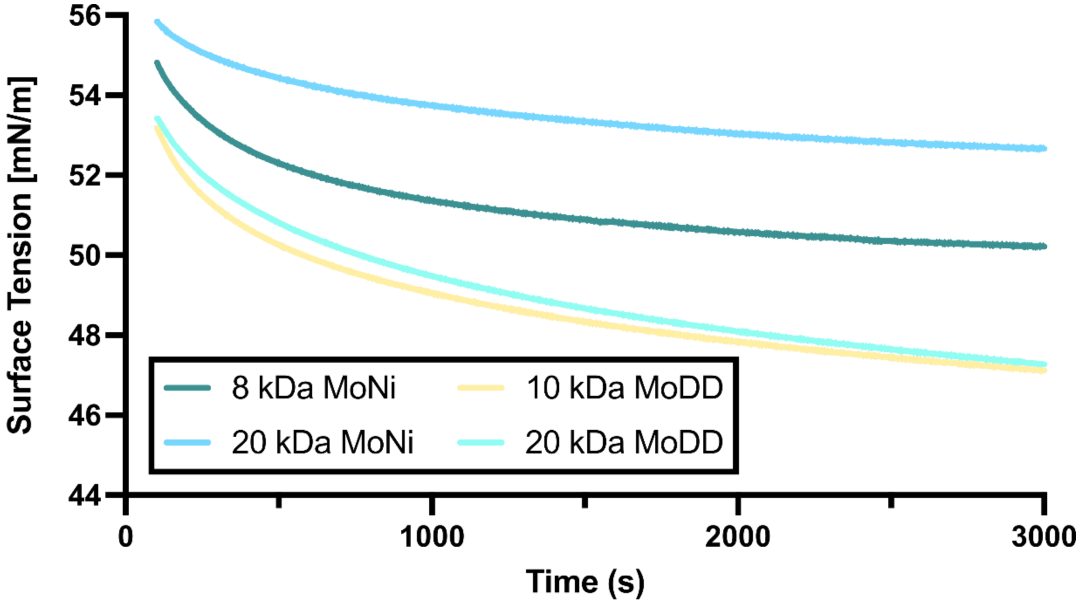

**Figure S23.** Evaluation of time dependent surface tension with Wilhelmy plate method for MoDD polymers compared to MoNi V70 variants. Mean surface tension values of 0.02 wt % polymer in Humalog buffer (without insulin) (n = 2).

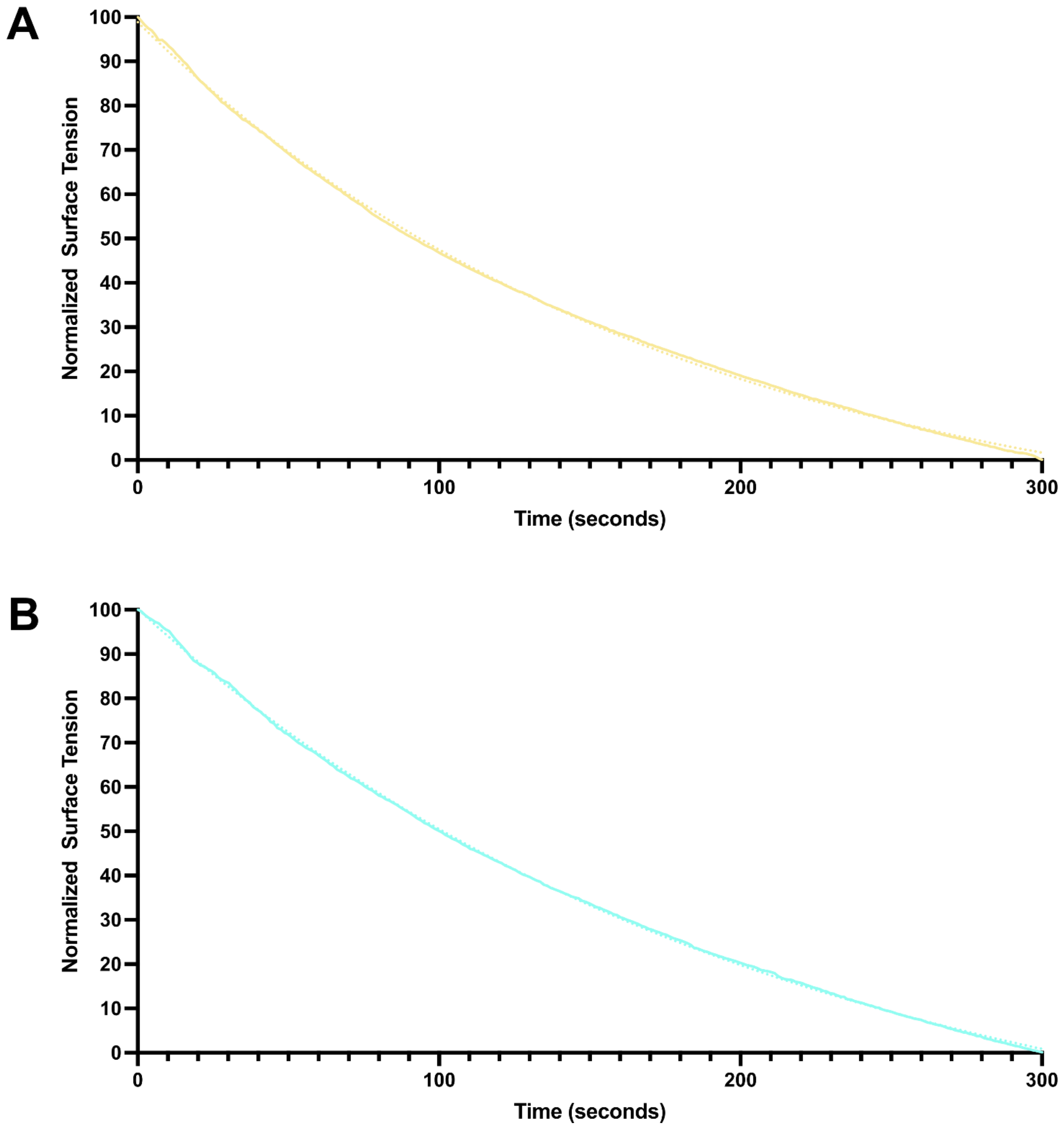

**Figure S24.** A) Curve fitting of the first 300 s (5 min) of time-resolved surface tensions measurements for 0.02 wt % solutions of MoDD10 in simulated Humalog buffer. B) Curve fitting of the first 300 s (5 min) of time-resolved surface tensions measurements for 0.02 wt % solutions of MoDD20 in simulated Humalog buffer. Experimental data is presented as the solid line and fits as the dotted lines. All fits were performed with Prism 10.

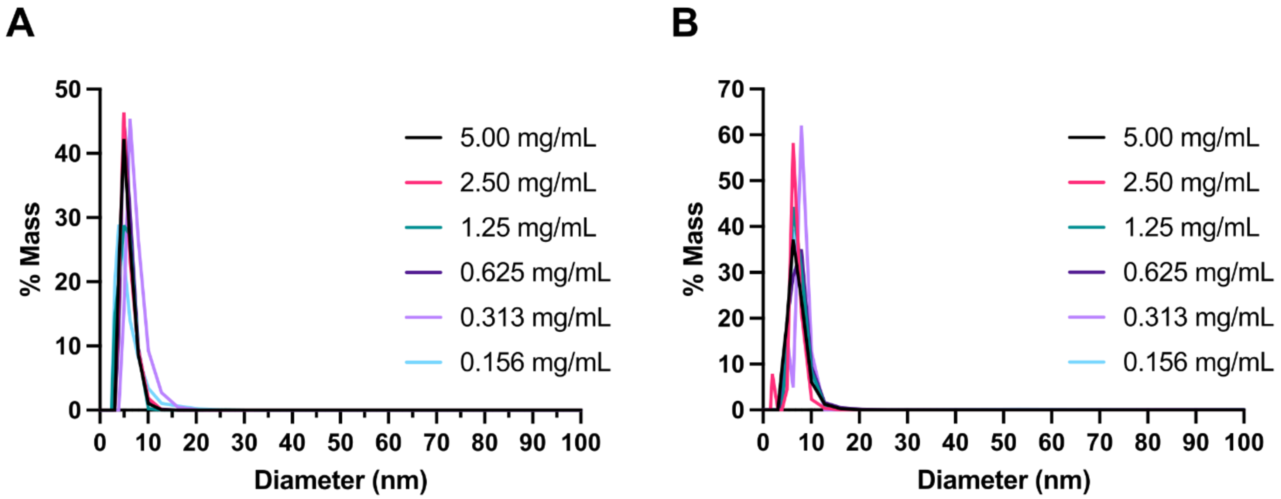

**Figure S25.** Dynamic light scattering measurements of MoDD polymers at various concentrations. No micellization was observed at the concentrations tested. A) MoDD10 B) MoDD20

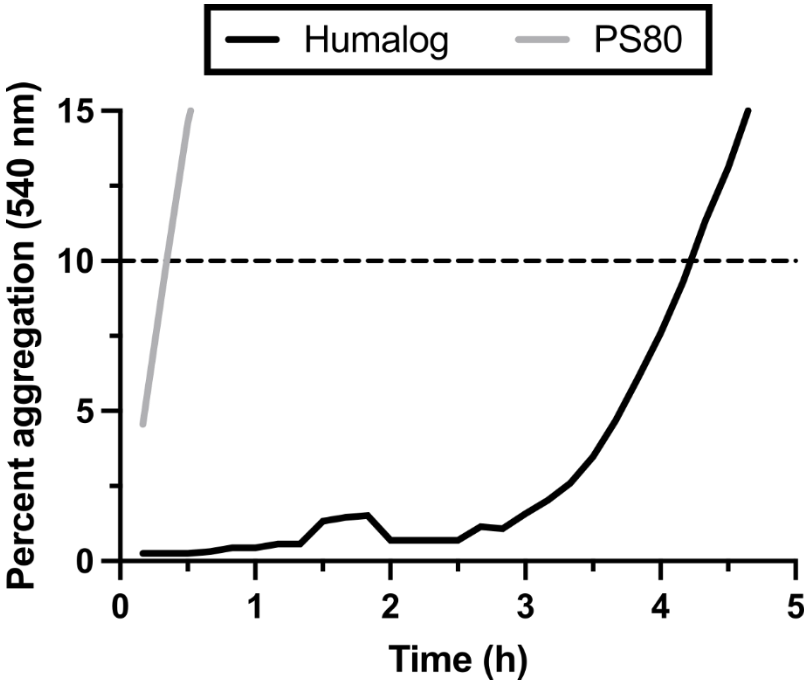

**Figure S26.** Non-specific absorbance assay to assess stability under stressed aging conditions (37 ˚C and shaking) of commercial Humalog and monomeric insulin with EDTA (1.2 mM) + PS80 (0.02 wt %). Data presented are a mean of 4 technical replicates

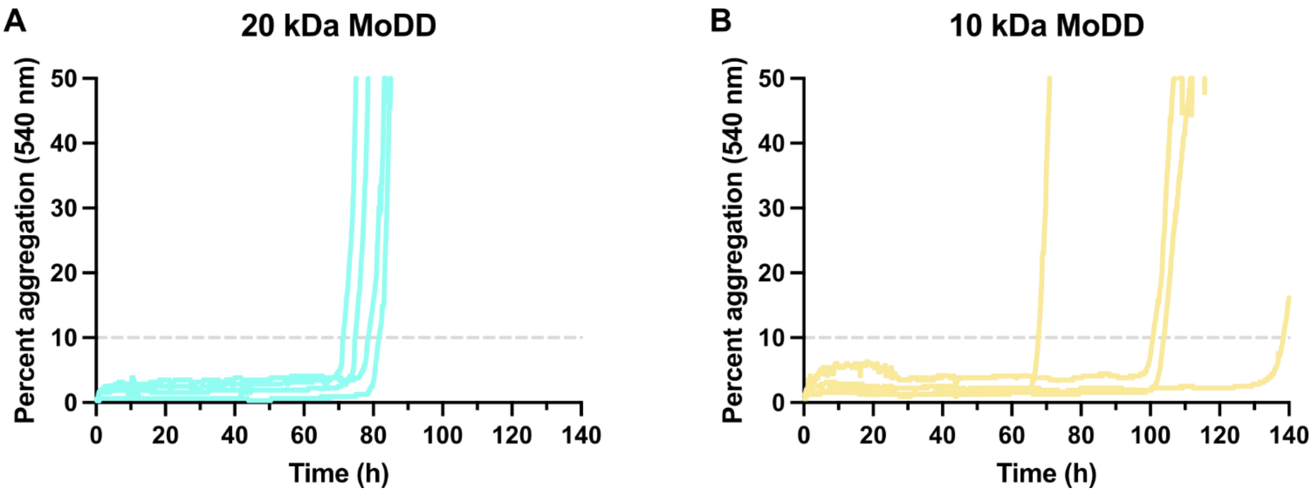

**Figure S27.** Non-specific absorbance assay to assess stability under stressed aging conditions (37 ˚C and shaking). Representative non-specific absorbance assays (n=4) of monomeric insulin with EDTA (1.2 mM) stabilized with A) MoDD20 and B) MoDD10.

**Table S2.** Summary of MoDD characteristics as a function of molecular weight. $M_n$* determined by conversion with [1]H NMR. $M_n$ and $M_w$ determined via SEC with polymethylmethacrylate (PMMA) standards. Equilibrium surface tension means and standard errors for duplicate measurements of 0.02 wt % solutions of MoDD polymers in Humalog Buffer as measured by the Wilhelmy plate method. Rate constants from one phase exponential decay of surface tension measurements in the first 5 min of equilibration. Compositional Dispersity Index (CDI) for each polymer determined from Compositional Drift simulations. Summary of T10 aggregation times for each polymer evaluated by insulin stability assay (n = 5).

| Property | 10 kDa MoDD | 20 kDa MoDD |
|---|---|---|
| $M_n$* | 10,300 | 20,600 |
| $M_n$ | 5,600 | 16,600 |
| $M_w$ | 7,061 | 19,800 |
| Dispersity | 1.26 | 1.19 |
| Equilibrium Surface Tension (mN/m) | 45.8 ± 0.3 | 45.8 ± 0.4 |
| Surface Kinetics Rate constant (k $10^{-3}$ $s^{-1}$) | 5.642 ± 0.130 | 4.827 ± 0.052 |
| Compositional Dispersity Index | 1.64 | 2.02 |
| 10% Aggregation Time (h) | 103.00 ± 11.40 | 77.63 ± 1.85 |

#### References

(1) Chen, M.; Moad, G.; Rizzardo, E. Thiocarbonylthio End Group Removal from RAFT-Synthesized Polymers by a Radical-Induced Process. Journal of Polymer Science Part A: Polymer Chemistry **2009**, 47 (23), 6704–6714. https://doi.org/10.1002/pola.23711.

(2) Mann, J. L.; Maikawa, C. L.; Smith, A. A. A.; Grosskopf, A. K.; Baker, S. W.; Roth, G. A.; Meis, C. M.; Gale, E. C.; Liong, C. S.; Correa, S.; Chan, D.; Stapleton, L. M.; Yu, A. C.; Muir, B.; Howard, S.; Postma, A.; Appel, E. A. An Ultrafast Insulin Formulation Enabled by High-Throughput Screening of Engineered Polymeric Excipients. Sci. Transl. Med. **2020**, 12 (550), eaba6676. https://doi.org/10.1126/scitranslmed.aba6676.

(3) Webber, M. J.; Appel, E. A.; Vinciguerra, B.; Cortinas, A. B.; Thapa, L. S.; Jhunjhunwala, S.; Isaacs, L.; Langer, R.; Anderson, D. G. Supramolecular PEGylation of Biopharmaceuticals. Proc Natl Acad Sci U S A **2016**, 113 (50), 14189–14194. https://doi.org/10.1073/pnas.1616639113.

(4) Maikawa, C. L.; Mann, J. L.; Kannan, A.; Meis, C. M.; Grosskopf, A. K.; Ou, B. S.; Autzen, A. A. A.; Fuller, G. G.; Maahs, D. M.; Appel, E. A. Engineering Insulin Cold Chain Resilience to Improve Global Access. Biomacromolecules **2021**, 22 (8), 3386–3395. https://doi.org/10.1021/acs.biomac.1c00474.

(5) Klich, J. H.; Kasse, C. M.; Mann, J. L.; Huang, Y.; d'Aquino, A. I.; Grosskopf, A. K.; Baillet, J.; Fuller, G. G.; Appel, E. A. Stable High-Concentration Monoclonal Antibody Formulations Enabled by an Amphiphilic Copolymer Excipient. Advanced Therapeutics **2023**, 6 (1), 2200102. https://doi.org/10.1002/adtp.202200102.

(6) Yu, H.; Liu, L.; Yin, R.; Jayapurna, I.; Wang, R.; Xu, T. Mapping Composition Evolution through Synthesis, Purification, and Depolymerization of Random Heteropolymers. J. Am. Chem. Soc. **2024**, 146 (9), 6178–6188. https://doi.org/10.1021/jacs.3c13909.

(7) Dykeman-Bermingham, P. A.; Bogen, M. P.; Chittari, S. S.; Grizzard, S. F.; Knight, A. S. Tailoring Hierarchical Structure and Rare Earth Affinity of Compositionally Identical Polymers via Sequence Control. J. Am. Chem. Soc. **2024**, jacs.4c00440. https://doi.org/10.1021/jacs.4c00440.
